# An mRNA SARS-CoV-2 vaccine employing Charge-Altering Releasable Transporters with a TLR-9 agonist induces neutralizing antibodies and T cell memory

**DOI:** 10.1101/2021.04.14.439891

**Authors:** Ole A.W. Haabeth, Julian J.K. Lohmeyer, Adrienne Sallets, Timothy R. Blake, Idit Sagiv-Barfi, Debra K. Czerwinski, Blaine McCarthy, Abigail E. Powell, Paul A. Wender, Robert M. Waymouth, Ronald Levy

## Abstract

The SARS-CoV-2 pandemic has necessitated the rapid development of prophylactic vaccines. Two mRNA vaccines have been approved for emergency use by the FDA and have demonstrated extraordinary effectiveness. The success of these mRNA vaccines establishes the speed of development and therapeutic potential of mRNA. These authorized vaccines encode full-length versions of the SARS-CoV-2 spike protein. They are formulated with Lipid Nanoparticle (LNP) delivery vehicles that have inherent immunostimulatory properties. Different vaccination strategies and alternative mRNA delivery vehicles would be desirable to ensure flexibility of future generations of SARS-CoV-2 vaccines and the development of mRNA vaccines in general.

Here, we report on the development of an alternative mRNA vaccine approach using a delivery vehicle called Charge-Altering Releasable Transporters (CARTs). Using these inherently nonimmunogenic vehicles we can tailor the vaccine immunogenicity by inclusion of co-formulated adjuvants such as oligodeoxynucleotides with CpG motifs (CpG-ODN). Mice vaccinated with the mRNA-CART vaccine developed therapeutically relevant levels of RBD-specific neutralizing antibodies in both the circulation and in the lung bronchial fluids. In addition, vaccination elicited strong and long lasting RBD-specific T_H_1 T cell responses including CD4^+^ and CD8^+^ T cell memory.

## Introduction

Coronavirus pandemics have been a growing concern for more than a decade and several attempts have been made to develop vaccines against SARS-CoV-1 and the Middle Eastern Respiratory Syndrome (MERS).^1^ The global spread of the SARS-CoV-2 virus stimulated worldwide efforts to leverage previous insights from coronavirus vaccines to develop safe, effective and scalable vaccines to alleviate the COVID-19 pandemic. While a variety of vaccine candidates and approaches are being investigated worldwide,^2^ the extraordinary pace of development and implementation of mRNA vaccines^3–6^ illustrates the potential of this emerging technology. The mRNA vaccines granted emergency use authorization by the FDA against SARS-Cov-2 represents a triumph of basic and applied science as these advances enabled the most rapid clinical translation from concept to clinical trial ever for a vaccine.^5,6^ mRNA is transiently expressed, does not integrate into the genome and is eliminated through natural degradation mechanisms in the body. mRNA vaccines offer flexible and fast design, that will allow for subsequent generations of products to address the emergence of new virus variants. The currently approved mRNA vaccines^5,6^ generated by in vitro transcription use chemically-modified nucleotides incorporated in mRNAs encoding the full viral spike protein, usually containing 2 structural epitope mutations, formulated in lipid nanoparticles (LNPs) and are administered intramuscularly. Despite their extraordinary success, the underlying science that contributes to the most effective, safe, and scalable vaccine against COVID-19 continues to evolve. Modifications and sequence optimization of the mRNA and the particular components of the delivery vehicle can influence the immunological response.^7^ Previous studies of SARS-CoV and MERS have shown that the proper choice of encoded antigen is critical^8^ to avoid potential complications from antibody dependent enhancement (ADE) of disease.^9^ The chemistry of the delivery vehicle is also important as the ionizable lipids that are a component of LNPs act as adjuvants but can induce adverse events^10^ and the use of polyethylene glycol (PEG) in the LNP formulations can contribute to allergic reactions.^11,12^

These continuing challenges and the degree to which the global scale of the COVID-19 pandemic has strained supply chains for the existing LNP technologies highlight the need to develop alternative approaches, both to enhance the resilience of the global response in the face of this and future pandemics, as well as to test alternative approaches that might lead to the most effective and durable immunological responses as well as to be able to respond rapidly to emerging new virus variants^13^.

In this study we present an alternative 3-component mRNA vaccine utilizing mRNA encoding the receptor binding domain (RBD) of the SARS-CoV-2 spike protein formulated with a highly efficient, non-toxic, PEG-free mRNA delivery platform called Charge-Altering Releasable Transporters (CARTs)^14–16^ and a TLR9 agonist (CpG) as co-formulated adjuvant^16,17^ (Fig. 1A). Prior studies had indicated that the RBD is a promising antigen target^18–20^ as antibodies elicited against RBD are often strongly neutralizing. CARTs offer a promising new gene delivery platform and have proven to be effective deliverers of mRNA vaccines in preclinical mouse studies.^16,21^ CARTs are single component amphiphilic diblock oligomers containing a sequence of lipid monomers and a sequence of cationic monomers (Fig. 1B, supplementary Fig. 1A). They are readily produced on scale in a two-step organocatalytic oligomerization. CARTs electrostatically encapsulate mRNA (or other co-formulated nucleotides like CpG) and deliver the genetic cargo into cells (supplementary Fig. 1B). A unique feature of CARTs is their ability to undergo a charge-altering rearrangement to produce neutral diketopiperazine small molecules (DKPs). This transformation facilitates the release of mRNA (Fig. 1C) and eliminates any toxic issues associated with persistent cations.^22^ Previous observations had shown that upon intravenous (IV) injection, CARTs containing hydroxyethyl glycine repeating cation units selectively deliver mRNA to the spleen,^14–16^ whereas intramuscular (IM) injections of these same products result in mRNA translation locally in the muscle (Fig. 1D). Bioluminescence studies with fLuc mRNA indicate in vivo protein expression is greater in the spleen after IV injection than in the muscle after IM injection. In either site, expression peaks after 4-6 hours (Fig. 1E) and decreases over a period of 3-4 days. Moreover, CARTs containing unsaturated lipid blocks exhibit enhanced transfection of antigen presenting cells^15^ motivating our choice of such a CART for the COVID-19 vaccine. The CART delivery vehicle does not induce nonspecific immune stimulation by itself.^16,21^ This property allows the co-formulation of oligodeoxynucleotide adjuvants such as the TLR9 agonist CpG-ODN to tune the induced immune response.

**Figure 1).**
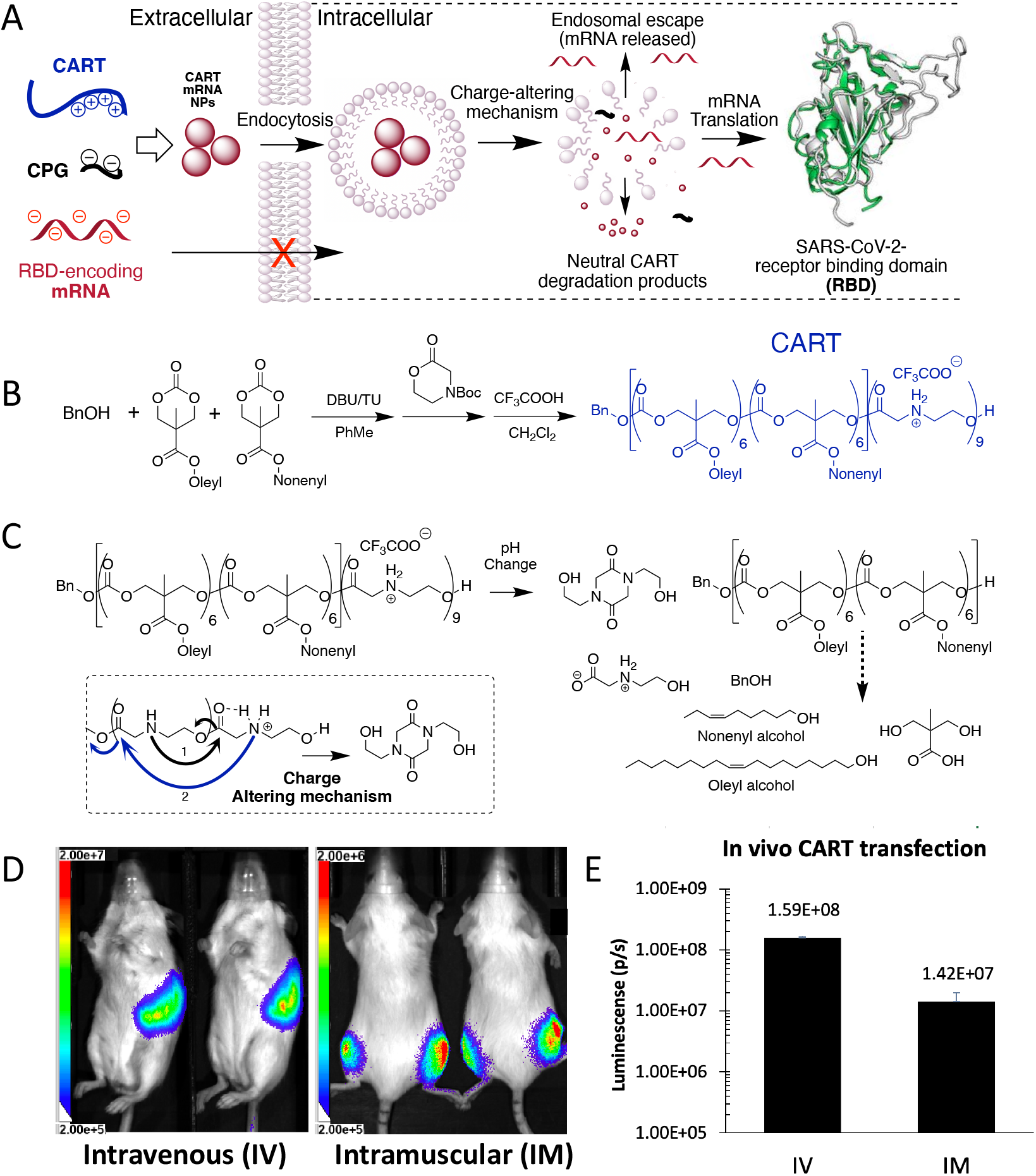
CART delivery platform methodology effectively complexes, delivers and releases mRNA via both systemic and local administration. (A) CART electrostatic formulation, cellular uptake, endosomal escape, and translation of SARS-CoV-2 RBD mRNA (B) CART synthesis via ring-opening polymerization (DBU=1,8-Diazabicyclo[5.4.0]undec-7-ene, TU=1-(3,5-bis(trifluoromethyl)phenyl)-3-cyclohexylthiourea) (C) CART chemical structure, degradation products, and charge-altering mechanism (D) In vivo luciferase reporter gene expression via systemic IV administration (left, 5μg fLuc mRNA), and local IM administration (2.5μg fLuc mRNA each flank) (E) quantification of in vivo mRNA expression at 4 hours post administration

Here we show that co-formulation of a CART with mRNA encoding SARS-CoV-2 RBD with the TLR9 agonist CpG (RBD mRNA + CpG-CART) induces robust neutralizing antibodies and RBD-specific T cell responses in mice. Moreover, we detect significant levels of these antibodies and memory T cells in the spleen and lung of vaccinated animals.

## Results

### RBD mRNA + CpG-CART vaccination elicits anti-RBD specific antibodies at day 4 after immunization

An mRNA encoding the Receptor Binding Domain (RBD)^23^ of SARS-CoV-2 was made by in vitro transcription and based on the published sequence of the virus. A His tag was included to allow for protein detection of the translated mRNA product as a quality control step. The resulting mRNA contained the optimal CAP-1 structure, uridines were replaced with modified N1-methyl-pseudouridine and cytidines were replaced with 5-methylcytidine to maximize mRNA stability and translation. Protein expression was verified by Western Blot in transfected 293F and Hela cells (supplementary Fig. 2A). To demonstrate that administration of mRNA-CART complexes does not lead to nonspecific immune stimulatory effects when formulated with optimally modified mRNA^24^, mice were injected intravenously (IV) with mRNA-CART and monitored for the activation of innate immune cells, a test that we have found to be most sensitive. mRNA-CART complexes were free of such nonspecific stimulatory effects when formulated with modified mRNA. However, when formulated with unmodified mRNA, nonspecific activation of innate immune cell subsets could be observed, demonstrating that the property of nonspecific immune stimulation is dependent on the cargo and not the CARTs (Supp. Fig. 2B). We were then able to direct the immune response of our candidate vaccine by co-formulating the mRNA-CART complexes with CpG oligonucleotides that trigger endosomal TLR9 receptors in antigen presenting cells. These CpG entities are known to be safe and effective vaccine adjuvants.^17^

We assessed the effects of the RBD mRNA-CART vaccination with or without the addition of CpG as adjuvant. IV injection of CARTs effectively delivers the cargo to antigen presenting cells (APCs) such as B cells, macrophages and dendritic cells in the spleen^15,16^ (Fig. 1D). This route of administration of mRNA-CART vaccination had proven effective in previous studies of therapeutic cancer vaccination.^16^ Based on this knowledge, we evaluated our vaccine first by IV administration. Mice were primed on day 0 and received two boosts on day 4 and on day 8 with 3ug RBD mRNA-CART formulated with or without 3ug of CpG (Fig. 2A). As control, a group that received 3ug CpG-CART alone with no mRNA and a control group treated with Phosphate Buffered Saline (PBS) were included. Mice vaccinated with RBD mRNA-CART plus CpG developed detectable levels of anti-RBD IgG and IgM as early as 4 days after vaccination (supplementary Fig. 3A). Over time, we observed an increase in the levels of anti-RBD antibodies in the serum of both mRNA-treated groups compared to controls (Fig. 2B). Importantly, the antibodies induced by our vaccine were specific for RBD and did not cross react to an irrelevant His tagged protein (GFP-His) (supplementary Fig. 3B). Notably, RBD-specific antibody responses in mice vaccinated with the formulation including CpG consistently exceeded those observed in groups treated without CpG (Fig. 2B, D).

**Figure 2).**
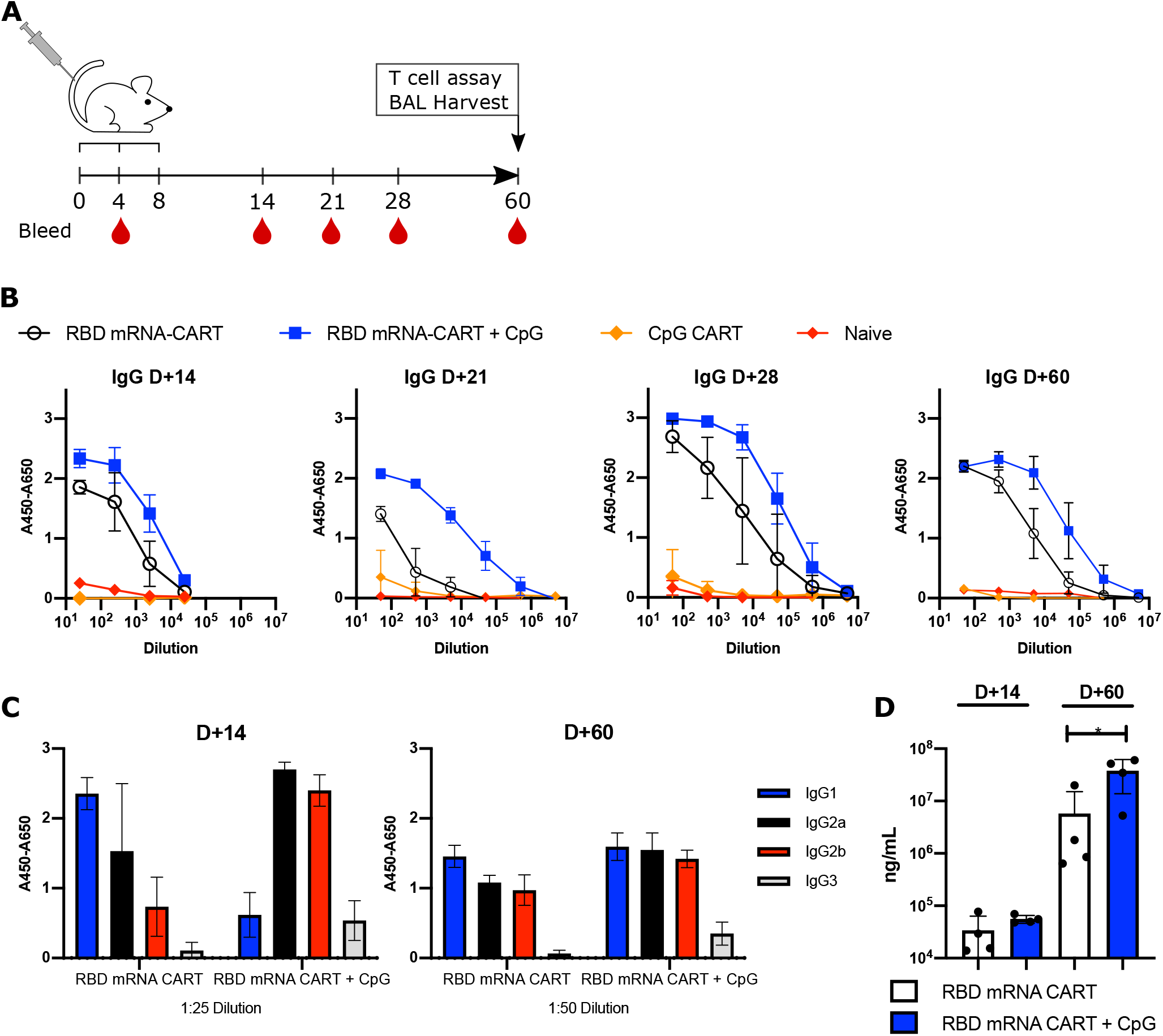
Addition of CpG to RBD mRNA-CART elicits a stronger anti-RBD immunoglobulin response and leads to earlier isotype switching. BALB/c mice were immunized intravenously with either 3ug RBD mRNA-CART (n=5), 3ug RBD mRNA-CART plus 3ug CpG (n=5), 3μg CpG CART (n=5) or Naïve untreated (n=5) and boosted on Day 4 and Day 8 after priming (A). Serum levels of RBD-specific IgGs from RBD mRNA-CART (Black), RBD mRNA + CpG-CART (Blue), CpG CART (Orange), and Naïve (Red) mice were monitored over the course of 60 days post priming by ELISA (B). On Day 14 and Day 60 after priming the distribution of IgG isotypes specific to RBD was analyzed using anti-mouse IgG1 (Blue), IgG2a (Black), IgG2b (red) and IgG3 (grey) monoclonal antibodies by ELISA (C). *On Day 14 and Day 60 after priming absolute concentration of the anti-RBD IgG was evaluated (D). *= P<0.05 unpaired student t-test (two-tailed)*. Data representative of 2 individual experiments.

### Addition of CpG results in early isotype switching of RBD-specific antibody responses

By day 14, we observed that the CpG co-formulated vaccine induced an RBD-specific Ig response that had undergone isotype switching from IgG1 to IgG2a, IgG2b and IgG3. By contrast, mice receiving the vaccine formulation without co-formulated CpG produced predominantly unswitched IgG1 (Fig. 2C and supplementary Fig. 4A, B). By day 60, these differences were less apparent, however we detected higher levels of all classes of RBD specific antibodies in serum of mice vaccinated with the CpG-containing vaccine (Fig. 2C and supplementary Fig. 4 B).

### Antibodies induced by RBD mRNA + CpG-CART vaccination inhibit RBD-ACE2 binding and neutralize pseudotyped viral entry

The induced anti-sera were next tested for their ability to inhibit the RBD-ACE2 interaction (Fig. 3A) and to block pseudotyped viral entry into ACE2 expressing target cells (Fig. 3B). Functional RBD-ACE2 receptor blocking was assayed both against the binding of RBD protein to the ACE2 expressing target cell by flow cytometry (data not shown) and by binding to solid phase coated with ACE2 protein. On day 28, we observed a striking difference in the level of neutralizing antibodies between the group that had received the vaccine containing the CpG as compared to the group vaccinated without the adjuvant. While anti-sera from RBD mRNA + CpG-CART vaccinated mice displayed a high degree of receptor blocking and pseudotyped viral neutralization with IC50 of 1:500 and 1:18000 respectively, the anti-sera from mice vaccinated with RBD mRNA-CART was only barely positive above background (Fig. 3A, B). However, by day 60 antisera from mice vaccinated with RBD mRNA-CART were able to block RBD-ACE2 receptor binding and neutralize pseudotyped viral entry, although to a lesser extent than antisera from mice vaccinated with RBD mRNA + CpG-CART (Fig. 3A, B). The results from the receptor interaction blocking assay and the pseudovirus neutralization assay were well correlated.

**Figure 3).**
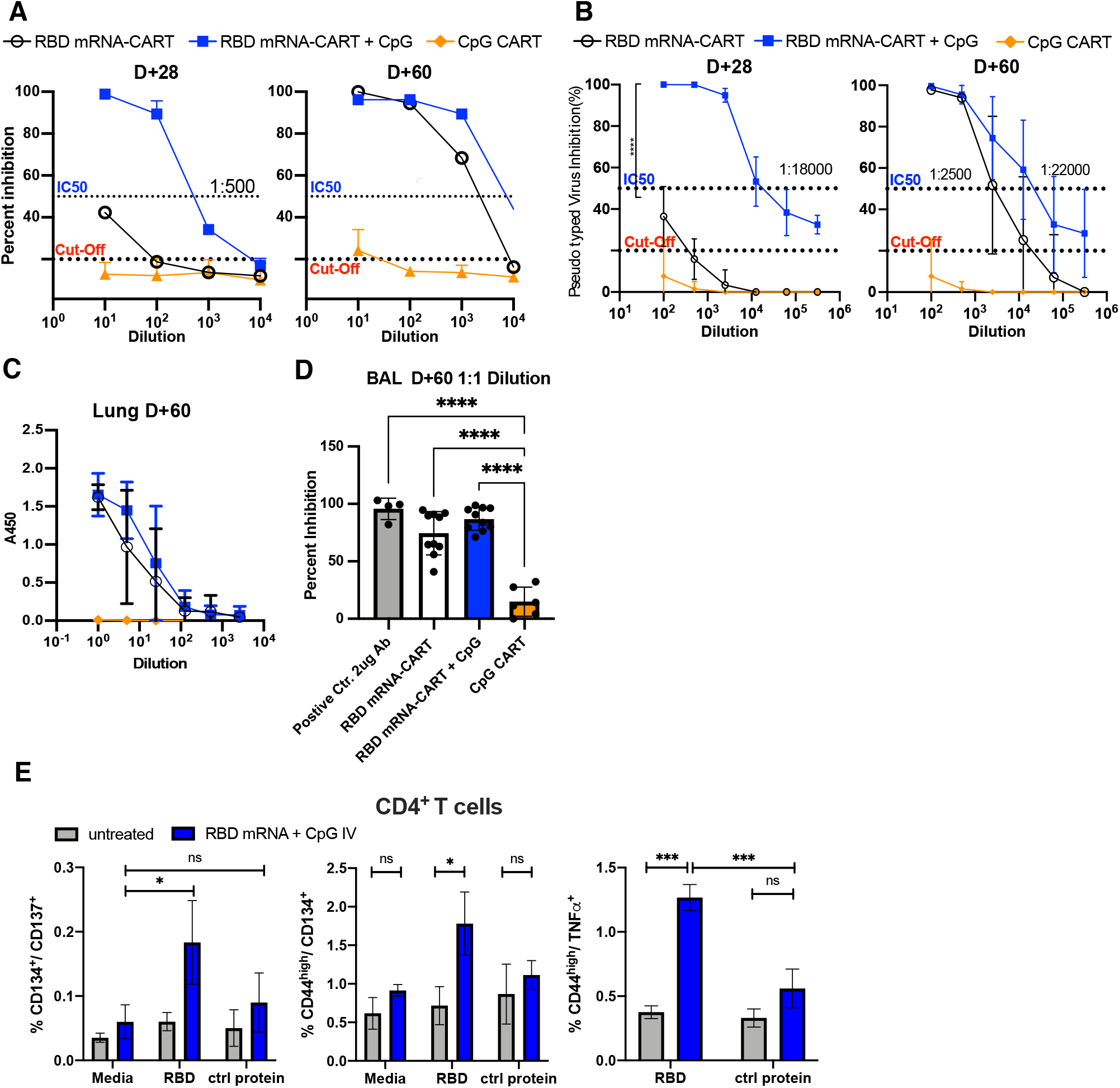
RBD mRNA + CpG-CART generates early high levels of RBD neutralizing antibodies. BALB/c mice (n=5) were immunized as described in Fig 2A. Sera from mice immunized with RBD mRNA-CART (Black), RBD mRNA + CpG-CART (Blue), CpG CART (Orange) were collected on D28 and D60 and tested in a commercially available RBD-ACE-2 inhibition assay (A). The same set of serum samples was tested in a pseudo typed virus neutralization assay. RBD expressing pseudo virus particles containing a zsGreen and Firefly Luciferase vectors were co-incubated with titrated concentrations of heat inactivated mouse serum. The pseudo virus particle-serum mix was then added to wells containing ACE-2 overexpressing 293F cells. Firefly luciferase expression was measured at 48h and 72h post experiment start (B). On D60 BAL was harvested from RBD mRNA-CART (Black), RBD mRNA + CpG-CART (Blue) and CpG-CART (Orange) immunized mice. RBD-specific total IgG was assayed by ELISA (C). BAL containing immunoglobulins were tested for their ability to inhibit binding of RBD to hACE-2 using a commercial ACE-2 inhibition kit (D). Lung single cell suspensions from naive mice (grey, n=3) or mice vaccinated on D0 and D21 IV with 3μg RBD-mRNA + 3μg CpG (blue, n=3) were collected on D28 and incubated with media alone, RBD protein or an irrelevant protein (CD81-His) [5μg/ml] for 48h and stained for T cell activation markers CD134, CD137 and intracellular TNFα on CD4^+^ T cells (E). Data are shown as mean ± SD. Data representative of 3 independent experiments (B-D) and 1 experiment (E). *= P<0.05, **=P<0.01 ***=P<0.001, ****=P<0,0001 one-way ANOVA (Tukey’s multiple comparison test) (D) or two-way ANOVA (Tukey’s multiple comparison test) (E)

### RBD-specific antibodies in bronchoalveolar lavage of vaccinated mice

To evaluate the presence of RBD-specific immunoglobulin in the lungs of vaccinated mice, we collected bronchoalveolar lavage (BAL) on D60 after treatment. RBD-specific Ig was detected in BAL from both RBD mRNA + CpG-CART and RBD mRNA-CART vaccinated mice by ELISA (Fig. 3C). Notably, as BAL is collected by flushing lungs with 2ml sterile PBS, the Ig titers represent highly diluted samples. Importantly, although diluted, these immunoglobulins from BAL of both vaccinated groups blocked RBD-ACE2 binding on D60 (Fig. 3D).

### RBD specific CD4^+^ T cells in the lung of vaccinated mice

In mice, Ig class switching is linked to T_H_1 T cell responses.^25,26^ To evaluate the vaccine induced T cell responses we prepared single cell suspensions from the lungs of IV vaccinated mice on D28 and cultured the cells for 48 hours in media alone, in the presence of soluble RBD-His protein or a control protein (hCD81-His). Cells were then collected and assayed by flow cytometry for T cell activation using fluorochrome conjugated monoclonal antibodies for memory and activation-specific surface proteins and intracellular cytokines. Remarkably, upon RBD protein restimulation, a defined RBD-specific CD4^+^/CD44^high^/CD134^+^ and CD4^+^/CD44^high^/TNFα^+^ activated T cell subset could be identified in lung cell suspensions from mice vaccinated with RBD mRNA + CpG-CART that could not be detected when cells were cultured with media alone or in the presence of the control protein (Fig. 3E). TNF*α* secretion by CD4^+^ T cells is associated with a T_H_1 polarization.

### RBD mRNA + CpG-CART vaccine elicits robust anti-RBD Ig responses by intravenous, intramuscular and subcutaneous route of administration

To test the efficacy of our vaccine in relation to the route of administration (ROA), we compared RBD-specific Ig responses induced by immunizations given IV, IM, or SC. Mice were primed with 3ug RBD mRNA and 3ug CpG formulated in CARTs on day 0 and received a boost on day 8 (supplementary Fig. 5A). All routes of administration led to detectable neutralizing antibody titers at the analyzed timepoints D14 and D28, with no significant differences between IM and SC administration (supplementary Fig. 5B). There was a tendency towards higher titers in IV vaccinated mice. Isotype switching occurred independent of route of administration (supplementary Fig. 5C).

### RBD mRNA + CpG-CART vaccine induces robust RBD-specific Ig responses in a clinically relevant prime-boost regimen and is independent of CpG source

Guided by the immunization regimen chosen by the currently approved SARS-CoV-2 vaccines^27^, mice were primed with the vaccine on D0 and boosted on D21. Mice were immunized either IV or IM with 3ug RBD mRNA + CpG-CART (Fig. 4A). Confirming previous observations, robust responses were observed for both groups. IV immunized mice showed higher-statistically not significant-titers of anti-RBD antibodies in serum and in the BAL on both D21 and D28 (Fig. 4B-D). Anti-sera from both groups effectively inhibited RBD-ACE2 binding on D21 (supplementary Fig. 6B), although substantially more effective on D28 reflecting the difference observed in total RBD-specific IgGs between the two groups (Fig. 4C). Moreover, robust antibody responses against the complete spike protein were observed in both groups, although higher in the intravenous group (supplementary Fig. 6C). Thus, the vaccine induced anti-RBD response is primarily directed towards exposed RBD epitopes in the complete spike protein. Importantly, although CpG is required for robust isotype switched anti-RBD immunoglobulin, the response is independent of the source of CpG. We tested 4 different sources of CpG, three of the C-subclass of CpGs (CpG-C) and one of the B-subclass (CpG-B). No significant difference was observed between the different CpG-Cs (supplementary Fig. 6 and 7), while the B-subclass CpG underperformed compared to the CpG-Cs (data not shown).

**Figure 4).**
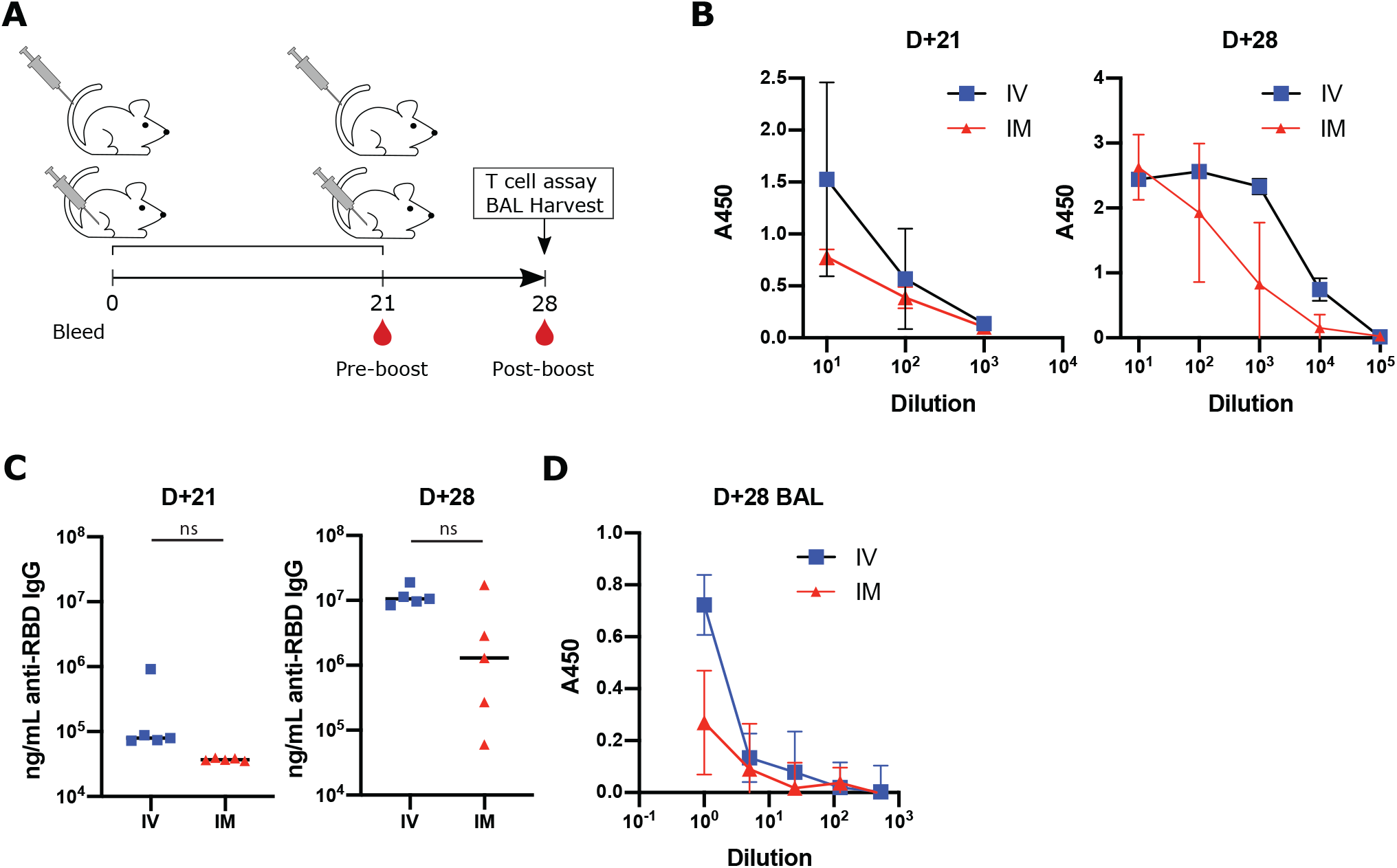
RBD mRNA + CpG-CART elicits neutralizing anti-RBD immunoglobulin responses after IV and IM vaccination. BALB/c mice (n= 5 per group) were immunized intravenously (IV) or intramuscular (IM) with 3ug RBD mRNA plus 3ug CpG and boosted on Day 21 after priming (A). RBD-specific immunoglobulin titers in serum were measured and quantified on Day 21 and Day 28 (B, C). On Day 28 BAL was harvested from both IV and IM treated mice and anti-RBD immunoglobulins were assayed by ELISA (D). Data are shown as mean ± SD. Data representative of 2 independent experiments. Statistical significance was assessed by Student’s t-test (two-tailed, unpaired) ns=P>0,05.

RBD mRNA + CpG-CART vaccine induced long lasting T_H_1 CD4^+^ and CD8^+^ T cell memory Splenocytes from mice that had received 3μg RBD-mRNA + 3μg CpG-CART or 3μg ctrl mRNA + 3μg CpG-CART either IV or IM on D1 and D21 were harvested on day 105 after vaccination and characterized for T cell responses by IFNγ enzyme-linked immunosorbent spot assay (ELISpot). In this assay, pooled splenocytes were enriched for either CD4^+^ or CD8^+^ T cells and cultured overnight with a SARS-CoV-2 RBD peptide pool or media alone. Significant IFNγ responses in CD4^+^ and CD8^+^ T cells were detected by both IV and IM vaccination. Since the route of administration of IV vaccinated mice targets the spleen, it is expected that spleen T cells would give a stronger response. On the other hand, it is remarkable to detect responding T cells 105 days after vaccination (Fig. 5 A, B).

**Figure 5).**
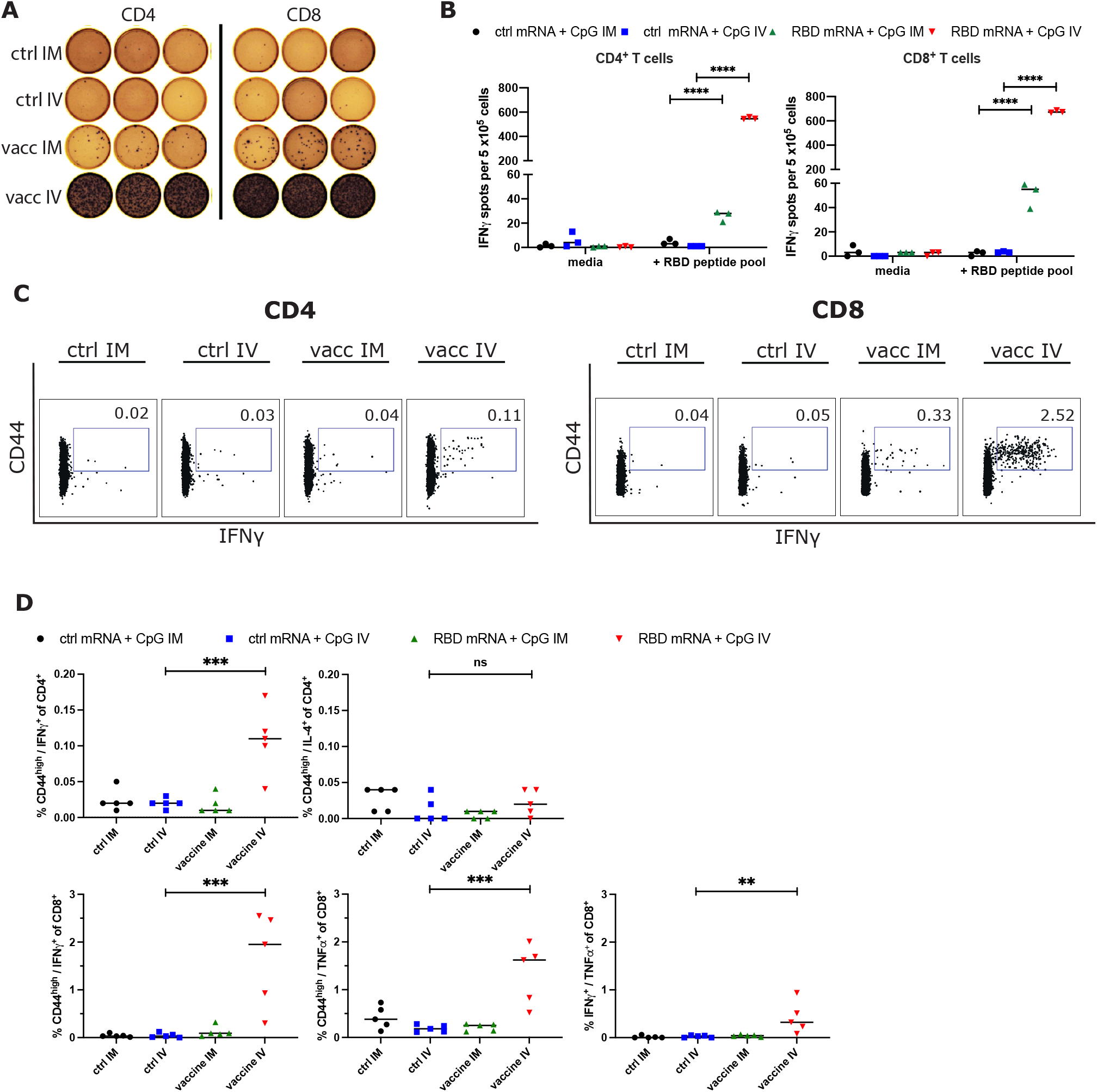
RBD mRNA plus CpG vaccination induces long lasting memory T_H_1 CD4^+^ and CD8^+^ T cell responses. BALB/c mice (n= 5 per group) were immunized intravenously (IV) or intramuscular (IM) with 3μg ctrl mRNA + 3μg CpG or 3μg RBD mRNA + 3μg CpG on D1 and boosted on D21 after priming. On day 105 pooled splenocytes were harvested, enriched for CD4^+^ and CD8^+^ T cells and stimulated separately for 16h with a RBD peptide mix for direct ex vivo IFNγ ELISpot assay (A, B). For ELISpot analysis splenocytes from the respective groups were measured in triplicates. Additionally, whole splenocytes of individual mice (n=5 per group) were incubated with media or a RBD peptide pool for 18h. After incubation cells were collected and stained for T cell memory marker CD44 as well as intracellular cytokines IFNγ, TNFα and IL-4 (C, D). Each dot represents the measurement of an individual mouse. **=P<0.01, ***=P<0.001, ****=P<0.0001 two-way ANOVA (B) or one-way ANOVA (D) (Tukey’s multiple comparisons test).

To further assess the functionality and polarization of the vaccine induced T cells we incubated splenocytes of individual mice from the same experiment in the presence of a RBD peptide pool or media alone. After 18 hours of incubation, CD4^+^ and CD8^+^ T cells were assayed separately by flow cytometry for their expression of memory markers CD44 and for the intracellular cytokines IFNγ, TNFα and IL-4. Even at this late time point after vaccination a significant population of RBD-specific IFNγ producing CD4^+^ and CD8^+^ T cells and TNFα producing CD8^+^ T cells could be identified in the RBD-mRNA + CpG-CART IV vaccinated group. There was no increase in IL-4 producing CD4^+^ T cells, indicating that T cell memory was predominately T_H_1 (Fig. 5 C, D, supplementary Fig. 8). In contrast to the IFNγ ELISpot results we were not able to identify these low frequency populations (mean 28 and 55 IFNγ spots per 5×10^5^ cells for CD4^+^ and CD8^+^ enriched conditions respectively) in IM vaccinated mice by flow cytometry.

### Neutralizing antibody levels of immunized mice are comparable to those achieved in vaccinated humans

Results from clinical trials indicate that the Pfizer/BioNTech mRNA vaccine and the Moderna vaccine can both confer protection from symptomatic infection prior to administration of their second booster vaccine doses.^28,29^ This implies that the antibody levels in humans at that early pre-boost time point are sufficient to confer disease protection. Accordingly, we compared the levels of neutralizing antibodies achieved in our vaccinated mice to those in immunized humans both prior to and after their boosters. BALB/c mice were vaccinated with 3μ RBD mRNA + 3μg CpG-CART either IM or IV on D1 and D21 and serum was collected on D28. These mice sera were then compared to sera from 13 individual Pfizer/BioNTech mRNA-LNP vaccinated humans collected 15-21 days after their priming vaccination and then again 15+/-4 days after their booster vaccinations. The level of RBD-ACE2 inhibition achieved with post boost sera from our IM and IV vaccinated mice was similar to or higher than that of the human pre-boost sera. The inhibitory antibody levels in the mice receiving the IV vaccination equaled those in humans post boosting (Fig. 6A, B).

**Figure 6).**
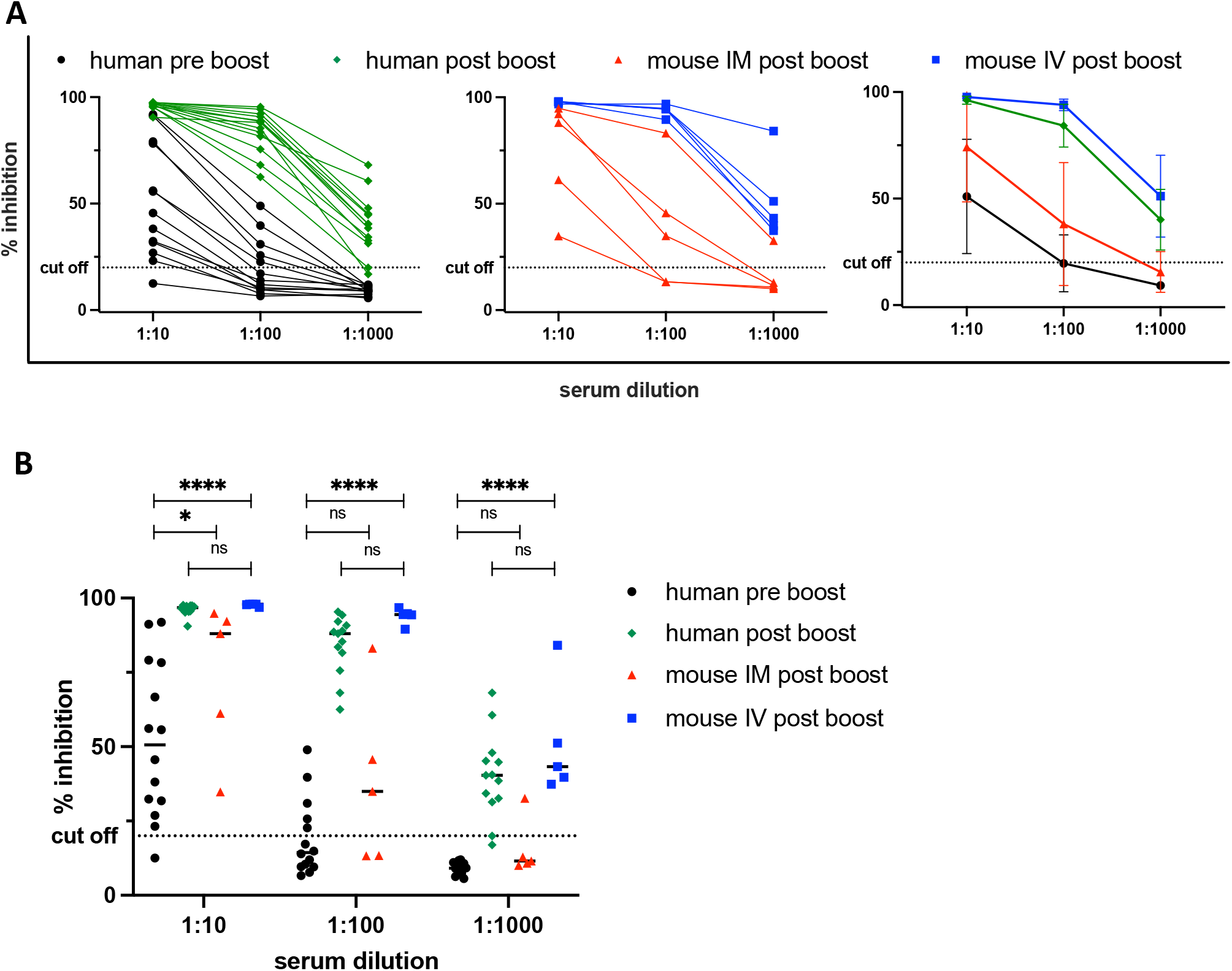
Neutralizing antibody levels of immunized mice are comparable to those achieved in vaccinated humans. Serum from immunized mice (IM in red, IV in blue, n=5) was harvested on Day 28. Serum from blood donors (n=13) who were vaccinated with the Pfizer/ BioNTech mRNA vaccine was collected either within 7 days before (pre boost, black) or 15±4 days after the boost (post boost, green) was tested for the ability to inhibit RBD/ACE-2 binding using a commercially available surrogate Virus Neutralization Test. *=P<0.05, ****=P<0,0001 one-way ANOVA (Tukey’s multiple comparison test)

### RBD mRNA + CpG-CART vaccination shows favorable safety profile

To evaluate the safety profile of the vaccine, mice were treated on D0 and D21 with either PBS, 3ug GFP mRNA-CART, 3ug GFP mRNA + 3ug CpG-CART or 3ug RBD mRNA + 3ug CpG-CART (supplementary Fig. S9A). No differences in bodyweight were observed after treatment (supplementary Fig. S9B, C). IV administration of CpG-containing formulations induced a transient decrease in White Blood Cell (WBC) count 24h after treatment that recovered by D2. This was driven by CpG, injection of mRNA-CART without CpG did not alter the WBC count (supplementary Fig. S9D- F). Serum levels of TNF*α* and IL-6 measured 1 day and 1 week after treatment were not affected by vaccination. IM and IV vaccination led to a transient increase of serum IP10 and IFN*α* levels 1 day after prime. Serum cytokine levels were within normal range at the second analyzed timepoint 7 days after prime. Again, transient effects were mediated by CpG, mice treated with mRNA-CART without CpG showed an unaltered cytokine profile after treatment compared to untreated mice (supplementary Fig. S9G- J). Similar dynamics for both WBC count and cytokine profile were observed after the boost treatment (data not shown). Treatment did not induce liver toxicity as assessed by serum liver enzyme levels of alanine transferase (ALT) and aspartate transferase AST (supplementary Fig. S9K- N) and alkaline phosphatase (AP) (data not shown). In addition, gross histopathology performed on day 1 and 5 after booster treatment revealed no pathologic findings in the gross assessment of the main organs.

## Discussion

For the first time mRNA-based therapeutics have been approved by the FDA and the success of both mRNA-based SARS-CoV-2 vaccines is both remarkable and mutually validating. However, challenges in production, deployment and availability of SARS-CoV-2 vaccines remain. One important aspect for the continuous success and refinement of mRNA-based therapeutics in general, will be to create access to diverse choices of safe delivery vehicles with varying chemical and biological properties. Here we demonstrate an effective mRNA vaccination strategy against the clinically relevant RBD antigen of SARS-CoV-2 using an alternative mRNA delivery platform to the clinically used lipid nanoparticles. We observed an effective SARS-CoV-2 specific immune response that was enhanced by the inclusion of the TLR9 agonist CpG as adjuvant. We further showed that the resulting anti-RBD sera were able to neutralize pseudoviral entry into ACE2-expressing cells. Focusing on the ACE2 receptor binding domain of the SARS-CoV-2 virus we were able to generate high amounts of isotype switched RBD-specific neutralizing antibodies in both serum and lungs of immunized animals. In addition, we compared serum levels of neutralizing antibodies from sera of humans vaccinated with approved Pfizer/BioNTech mRNA-LNP vaccines with those of mRNA-CART vaccinated mice. The levels of neutralizing antibodies in fully immunized mice were similar (IM) or higher (IV) than the levels of neutralizing antibodies measured in pre-boost blood samples from humans vaccinated with the Pfizer/BioNTech mRNA vaccine. Since it has been shown that this approved mRNA vaccine confers protection as early as day 12 post prime, this is an indication that our vaccine induced neutralizing antibody levels can be sufficient to confer protection.^28^ Of course, for further validation of these results vaccination studies in larger animals such as non-human primates are needed. Finally, the RBD mRNA + CpG-CART vaccine induced RBD-specific CD4^+^ and CD8^+^ T cell responses with induction of long-lasting T cell memory of T_H_1 polarization. In addition, the IgG isotype profile dominated by IgG2a and IgG2b confirms a T_H_1 polarized T cell response.

When compared to LNPs, CARTs have unique biodistribution, selectively delivering mRNA to the spleen or other organs without the need for targeting ligands, simply through changes in the CART structure. CARTs can be readily prepared and formulated with multiple mRNAs in any desired nucleotide combination^21^, only require a single structural component that is mixed with mRNA, and do not require specialized microfluidics instruments for their manufacture. This allows for alternative drug application strategies. Preliminary experiments show that CARTs formulated with mRNA are stable for 11 days at -20°C (supplementary Fig. S11). However, since formulation does not require specific equipment, a mix-and-shoot administration of the vaccine could be used. Our initial experiments indicate that unformulated CARTs in DMSO are stable for more than 12 months at -20°C. CARTs and mRNA could therefore be divided into two separate chambers of a two-chamber syringe, allowing for mixing and mRNA-CART formation at the point of administration and avoiding thermostability issues of pre-formulated complexes. In contrast to LNPs, CARTs have no unspecific immunostimulatory effects, which allows more flexibility for vaccine design and the option to vary oligodeoxynucleotide adjuvant quantity when co-delivered with mRNA, rather than relying on the inherent immunogenicity of LNPs. When CARTs are injected IM, gene expression localizes exclusively at the site of injection and does not spread to other organs. In a SARS-CoV-2 vaccine study using IM injection of LNPs, the liver showed the highest level of reporter gene and antigen expression.^19^ In addition, IV injection of CARTs confer mRNA expression exclusively to the spleen which could explain the higher potency of IV vaccination compared to IM. While IV administration of the vaccine induced a stronger antibody and T cell response, significant responses could be induced via both routes of administration. Interestingly, it has been shown recently that IV vaccination with a BCG vaccine against tuberculosis profoundly altered the protective outcome in non-human primates with an increase of antigen responsive CD4^+^ and CD8^+^ T cells in blood, spleen, BAL and lung lymph nodes when compared to the established intradermal or aerosol administration.^30^ Additionally, IV administered mRNA lipid nanoparticles have demonstrated potency in preclinical mouse models and a clinical phase I study of therapeutic cancer vaccination.^31,32^

We chose to direct our vaccine specifically against the RBD rather than the whole spike protein sequence. Potent humoral and cellular immune responses have been observed in clinical trials with both the SARS-CoV-2 full-length spike protein and RBD.^6^ Moreover, an RBD mRNA directed vaccine was proven safe in a phase 1/2 clinical trial tested in the US and in Germany.^27,33^ In addition, RBD provides essential targetability for humoral and cellular immune responses. Piccoli et al. showed that 90% of the neutralizing activity of serum from exposed patient target the RBD.^34^ RBD is also an epitope for T cell responses against SARS-CoV-2 S protein.^35^ A fast-spreading mutant with a mutation in the RBD (N501Y) has been identified in the UK (B.1.1.7) raising concerns about coverage of the current mRNA-based vaccines.^6^ This N501Y mutation does not seem to affect the efficacy of an RBD vaccination since mice vaccinated against the original RBD sequence were able to clear a SARS-CoV-2 variant containing the specific mutation.^36^ However recent data indicates that other clinically relevant RBD and non-RBD mutations can mediate escape from vaccine induced humoral immunity^13,37^, highlighting the urgent need of flexible and rapidly adaptable vaccine platforms.

The safety data of mRNA + CpG-CART vaccination seems to be favorable. The detected changes in white blood cell count and cytokine profile were mediated by CpG. However, CpG has a well-known safety record in clinical studies of other vaccines. We believe that the ability to formulate TLR activating molecules like CpG into our vaccine will aid in inducing a protective immune response in populations with less competent immune systems and that are more at risk for severe COVID19 symptoms. CpG directly activates pDCs and B cells, contributing to the induction of both innate and adaptive immune responses. The cascade of events initiated by CpG indirectly supports maturation, differentiation and proliferation of natural killer cells, T cells and monocytes/macrophages.^38–41^ B cells activated by CpG upregulate expression of their Fc receptor (FcR) and costimulatory molecules including MHC class II, CD40, CD80 and CD86.^42–44^ Subsequently, the CpG-stimulated B cells proliferate and differentiate into plasma cells and memory B cells.^45^ The adjuvant effects of CpG are supported by our study where the addition of CpG resulted in a more rapid immune response, higher anti-RBD titers in serum and bronchoalveolar lavage, more effective ACE2-RBD inhibition and pseudo typed virus neutralization, increased T cell response, and more pronounced isotype switching.

In conclusion, our study demonstrates the potency and flexibility of this mRNA-CART vaccine platform against the clinically relevant SARS-CoV-2 RBD antigen. The robust induction of both B and T cell responses via different routes of administration warrants further exploration and its use as an alternative to the clinically approved lipid nanoparticles in the general development of mRNA-based therapeutics against infectious diseases.

## Material and Methods

### Recombinant Proteins

The pCAGGS plasmids coding for soluble RBD-His, residues 319-541, and spike-His, residues 1-1213 from the Wuhan-Hu-1 genome sequence (GenBank MN9089473) were a gift from Prof. Florian Krammer (Icahn School of Medicine at Mount Sinai).^23^ The pcDNA3 plasmid coding for a soluble ACE2-hIgA FC fusion protein was purchased from addgene (ID 145154). Plasmids were expanded using One Shot™ TOP10 Chemically Competent E. coli (ThermoFisher Scientific) and the ZymoPURE II Plasmid Maxiprep Kit (Zymo Research). Recombinant proteins were produced using the Expi293F cells (Thermo Fisher Scientific) by transfecting 200×10^6^ of these cells with purified DNA using the ExpiFectamine 293 Transfection Kit (Thermo Fisher Scientific). Supernatants from transfected cells were harvested 3 days post transfection by centrifugation at 300g for 10 min and filtration through a 22um filter. RBD-His and Spike-his containing supernatants were batch purified using the HisPur™ Ni-NTA Resin (ThermoFisher Scientific). Supernatants were incubated with 6 ml of resin for 1h at RT. Then, the resin was recovered by centrifugation (2 min at 700g), washed, and finally eluted per manufacturer recommendation. Elution fractions were analyzed by SDS-PAGE and western blot to confirm RBD-His or Spike-His purification and the positive fractions were pooled.

### In vitro transcription

The RBD-6his was cloned from the pCAGGS expression plasmid into the LF-pLMCT plasmid that contains a T7 promoter and a polyA sequence required for mRNA synthesis. The LF-pLMCT plasmid was a gift from Dr. Kris Thielemans (Free University of Belgium). The RBD-6his coding sequence was amplified by PCR using PHUSION polymerase (NEB) and TAAACTTAAGACAACCATGGTCGTGTTTCTGGTGC as a forward primer and GGGGATCCcGTCTTCCTCGAGTTATCAATGGTGATGGTGA as reverse primer. The PCR product was then inserted into pLMCT by NcoI and XhoI. mRNA coding for RBD was synthesized per manufacturer recommendation using Hiscribe T7 (NEB) with co-transcriptionnal CleanCap AG (Trilink), N1-methyl-pseudouridine (Trilink) and 5-Methylcytidine (Trilink). The template for in vitro transcription was a PCR amplicon from the pLMCT-RBD-6His produced using the PHUSION high fidelity DNA polymerase (NEB) and TGTGGAATTGTGAGCGGATA as forward primer and CTTCACTATTGTCGACAAAAAAAAAAAAAAAAAAAAAAAAAAAAAAAAAAAAAAAAAAAAAAAAAAAAAAAAAAAAAAAAAAAAAAAAAAAAAAAAAAAAAAAAAAAAAAAAAAAAAAAAAAAAAAA as reverse primer.

### CART preparation and characterization

CART O_6_-*stat*-N_6_: A_9_, consisting of a first block of a 1:1 statistical mixture of oleyl and nonenyl-substituted carbonate monomers, followed by a block of α-amino ester monomer was prepared as previously reported.^14,15^ Briefly, to a mixture of nonenyl (29 mg, 1 mmol), and oleyl carbonate (40.5 mg, 1 mmol) in toluene (150 μL) was added TU, DBU, and BnOH (5 mol% TU/DBU, 0.2 mmol BnOH) in 50 μL toluene. The reaction was stirred for 1.5 hours, then the morpholinone monomer (33.4 mg, 0.16 mmol) was added as a solid then stirred for an additional 2.5 hours. The reaction was quenched with AcOH then dialyzed overnight in DCM/MeOH (3.5 kDa M.W, cut-off). Concentration after dialysis afforded 85 mg clear residue which was deprotected with TFA (0.85 mL) in dry DCM (8.5 mL) overnight. Endgroup analysis of the deprotected polymer showed block lengths of 6 nonenyl and 6 oleyl carbonate units and 9 cationic aminoester units.

### CART oligonucleotide formulation

To prepare the CART-vaccine, CARTs were formulated with a mixture of CpG and RBD mRNA at a 10:1 cation: anion ratio assuming full protonation of the CART and full deprotonation of the oligonucleotides (1:1 mass ratio of CpG and mRNA nucleotides). Formulations were prepared by mixing the reagents for 20 seconds in acidic PBS (pH adjusted to 5.5 by addition of 0.1M HCl) in a total volume of 50-100μl, followed by a brief spin in a tabletop centrifuge. The formulation was used within 5 minutes for in vitro or in vivo experiments.

### Mouse vaccination

Eight- to twelve-week-old female BALB/c mice were purchased from The Jackson Laboratory and housed in the Laboratory Animal Facility of the Stanford University Medical Center. All experiments were approved by the Stanford Administrative Panel on Laboratory Animal Care and were conducted in accordance with Stanford University Animal Facility and NIH guidelines.RBD-mRNA and CpG were formulated with CART polymer in PBS at pH 5.5 as described above. Mice were injected with 3μg RBD-mRNA formulated with 2.6μl CART (5mM) or 3μg CpG formulated with 2.6μl CART (5mM) or 3μg RBD-mRNA plus 3μg CpG formulated with 5.2μl of CART (5mM). Mice were vaccinated by IV, IM or SC injection and were boosted as described in the experiment. CARTs are formulated at indicated concentrations of mRNA in 50-100 μL total volume. For IV administration 100 μl formulated CART was administered per tail vein injection. For IM injections 50 μl of formulated CART was injected in the thigh muscle. SC injections were administered on the back of the mouse near the tail. At indicated time points mice were bled and serum was collected.

### Hela and 293F transfection

Hela cells and 293F cells were plated at 10^6^ cells per well in a 12 well plate in Opti-MEM media (ThermoFisher Scientific). 2 μg of the RBD-his mRNA or GFP mRNA (Trilink) was formulated in 6.6 μl of PBS pH 5.5 with 1.37 μl of 5mM CART and added to the cells. After 4 hours of transfection, Opti-MEM media was replaced by RPMI media containing 10 % FCS and Penicillin-Streptomycin 1000 U/ml. 12 hours post transfection RBD and GFP expression were monitored by western blot and fluorescence microscopy, respectively.

### Western blot

15ul of media from Hela or 293F transfected cells were mixed with 4x sample loading buffer (Invitrogen) and were loaded on a 4-12% NuPAGE gel (Invitrogen). Electrophoresis was performed in an MES buffer at 200V for 35 min. Proteins were transferred to a cellulose membrane using the iBLOT system (Invitrogen). The membrane was stained with Ponceau red to verify protein transfer then the membrane was blocked for 1h in TBST containing 5% non-fat dry milk. The membrane was washed 3 times in TBST and incubated in TBST containing 5% non-fat dry milk and 1:1000 mouse anti-His (Biolegend) overnight. After 3 washes in TBST the membrane was incubated with 1:10000 anti-mouse Ig (Southern Biotech) in TBST containing 5% non-fat dry milk for 1 hour. After 3 washes in TBST the blot was revealed using the EC™ Prime Western Blotting System (Sigma). The membrane was imaged using a Chemidoc MP imaging system from BioRad.

### Serum preparation

For human samples, informed consent was obtained from the subjects prior to blood draw. Blood was collected in Eppendorf tubes and allowed to coagulate for 60 min at room temperature. After 10 min of centrifugation at 1000xg the supernatant was collected. Serum was heat inactivated at 56°C for 30 min.

### ELISA

Nunc-Immuno™ MicroWell™ 96 well ELISA plates (MilliporeSigm) were coated overnight with 50ul per well of 2ug/ml RBD-His or Spike-His protein in carbonate buffer pH9. After 3 washes in ELISA wash buffer (PBS with 0.1% Tween 20), plates were blocked using 100ul of 5% nonfat dry milk diluted in TBS buffer containing 0.1% Tween 20 (TBST) for 1 hour at room temperature. Serum, BAL and antibody dilutions were prepared in TBST containing 1% nonfat dry milk. The blocking solution was removed and 50 μl of each serial dilution was added to the plate for 1 hour at room temperature. Plates were washed three times and incubated with HRP conjugated anti-human Ig (1:5000, BioSource), anti-mouse Ig (1:5000, Cell Signaling), anti-mouse IgG2a (1:5000, Southern Biotech), anti-mouse IgG2b (1:5000, Southern Biotech), anti-mouse IgG1 (1:5000, Southern Biotech), anti-mouse IgG3 (1:5000, Southern Biotech), anti-mouse IgA (1:5000, Invitrogene), or anti-mouse IgM (1:5000, Southern Biotech). Plates were washed three times and 100 μl of TMB ELISA Substrate (Abcam) were added to each well. ELISA was developed for 10 min and then the reaction was stopped by adding 50 μl of Stop Solution for TMB Substrates (ThermoFisher Scientific) to each well. In some assays, a human anti-RBD (Invivogene) of known antibody concentration was used as standard. Optical density at 450nm (OD450) was measured using a SpectraMax Paradigm Microplate Reader (Molecular devices).

### RBD-ACE2 interaction blocking assay ELISA

RBD-ACE2 interaction blocking assay was evaluated using three methods: a commercial kit from Genescript and an in house developed ELISA and by flow cytometry.

For the commercial kit, we used the SARS-CoV-2 Surrogate Virus Neutralization Test (sVNT) Kit (Genescript) following the manufacturer’s instructions. In short samples and controls were diluted at indicated ratios with Dilution buffer and pre-incubated with HRP-RBD in a 1:1 ratio for 30min at 37°C. Samples were then added to the capture plate in wells pre-coated with hACE2. After 15’ incubation at 37°C wells were washed four times with wash buffer. TMB solution was added and incubated for 15’ at room temperature in the dark. After 15’ stop solution was added to the wells and promptly analyzed. Optical density at 450nm (OD450) was measured using a SpectraMax Paradigm Microplate Reader (Molecular devices).

For the in-house developed ELISA, Nunc-Immuno™ MicroWell™ 96 well ELISA plates (Millipore) were coated overnight with 50 μl per well of 2ug/ml RBD-His or Spike-His protein in carbonate buffer pH9. After 3 washes in ELISA wash buffer (PBS with 0.1% Tween 20), plates were blocked using 100ul of 5% nonfat dry milk diluted in TBS buffer containing 0.1% Tween 20 (TBST) for 1 hour at room temperature. Serum, BAL and antibody dilutions were prepared in TBST containing 1% nonfat dry milk. The blocking solution was removed and 50 μl of each serial dilution was added to the plate for 1 hour at room temperature. Plates were washed three times and 50 μl of 2 times diluted ACE2-hIgA supernatant was added to each well for 1 hour. After 3 washes, plate was incubated with HRP conjugated anti-human IgA (1:1000, Thermo Scientific) for 1hour in TBST with 1% non-fat dry milk. Plates were washed three times and 100 μl of TMB ELISA Substrate (Abcam) were added to each well. ELISA was allowed to develop for 10 min and then the reaction was stopped by adding 50 μl of Stop Solution for TMB Substrates (ThermoFisher Scientific) to each well. In some assays human anti-RBD (Invivogen) of known antibody concentration was used as standard. Optical density at 450nm (OD450) was measured using a SpectraMax Paradigm Microplate Reader (Molecular devices).

For the flow cytometry assay, RBD-His 2ug/ml was incubated with sera for 1 hour. Then, the 4×10^5^ ACE2 expressing HEK293T cells were added to the RBD-His/sera mix and incubated at RT for 30 min. Cells were then washed 2 times in PBS containing 1% BSA. RBD was then detected using an Alexa Fluor 488 conjugated anti-His antibody (clone J099B1, Biolegend). Cells were analyzed on a flow cytometry (BD).

### Pseudo-virus assay

Pseudo-typed lentivirus expressing the Sars-Cov-2 spike protein and the luciferase was produced in HEK293T cells as previously described^46,47^. One day before transfection, 6 × 10^6^ HEK293T cells were seeded in a 10-cm culture plate in RPMI containing 10% FCS, 2mM L-glutamine, streptomycin and penicillin. Using TransIT (Mirus), cells were then transfected with 10 μg of the lentiviral packaging vector (pHAGE_Luc2_IRES_ZsGreen), the 3.4 μg of SARS-CoV-2 spike, and lentiviral helper plasmids (2.2 μg of HDM-Hgpm2, 2.2 μg of HDM-Tat1b, and 2.2 μg of pRC-CMV_Rev1b). The spike vector contained the full-length wild-type spike sequence from the Wuhan-Hu-1 strain of SARS-CoV-2 (GenBank NC_045512). These 5 plasmids were kindly provided by Dr. Jesse Bloom (Fred Hutch Seattle, University of Washington). 72 hours post transfection, virus-containing supernatant was harvested, centrifuged at 300g for 5 minutes, filtered on a 0.45 um filter, aliquoted and frozen at -80°C.

For viral neutralization assays, ACE2-expressing HEK293T^47^ cells were plated in poly-L-lysine-coated, white-walled clear-bottom 96-well plates at 12,500 cells/well 1 day prior to infection. Mouse serum was centrifuged at 2,000 x g for 15 min, heat inactivated for 30 min at 56 °C and diluted in D10 media (DMEM medium supplemented with 10% FCS). Virus was diluted in D10 medium, supplemented with polybrene (5 μg/mL), and then added to serum dilutions. The virus/serum mix was preincubated for 1 hour at 37°C before it was added to the cells and incubated at 37 °C for ∼48 h. Cells were lysed by adding BriteLite assay readout solution (Perkin Elmer) and luminescence values were measured with a SpectraMax Paradigm Microplate Reader (Molecular devices). As positive control a neutralizing human anti-SARS-Cov-2 IgG1 antibody was used (Acro).

### Bronchoalveolar lavage

Mice were sacrificed and lungs were inflated 2 times with 1 ml of PBS following a previously described procedure^48^. Lavage fractions were pooled and centrifuged at 1200 rpm for 5 min. Supernatant was collected and assayed for anti-RBD antibodies by ELISA.

### T cell response assay on lungs

Mouse lungs were harvested at indicated days after vaccination. To prepare lung single cell suspensions, lungs were cut into small pieces and incubated at 37°C in RPMI containing Collagenase D (2mg/ml, Sigma) and DNAse (50ug/ml, Sigma) for 30min. Then, digestion mix was diluted 5 times and the lung preparations were processed through a 70-um cell strainer. Red blood cells present in the spleen or lung single cell suspensions were lysed using ACK buffer (ThermoFisher Scientific). Single cell suspensions were kept on ice until further processing for T cell response assay. Cells were cultured in 96-well plates (Corning, V-bottom) at 1×10^6^ cells/well and stimulated for 48 hours with 5 μg/mL of RBD-His or hCD81-His or media alone (RPMI + 5% FCS) in the presence of anti-mouse CD28 antibody [0,5 μg/ml] (Southern Biotech). As positive control cells were stimulated with anti-mouse CD3 [0,05 μg/ml] (Southern Biotech). For intracellular staining, cells were treated with GolgiStop (BD Biosciences) for 5 hours prior to staining. Following stimulation, cells were washed, stained with Aqua Live/dead viability dye (Thermo Fisher) in PBS, washed two additional times and stained with a cocktail of monoclonal antibodies and Fc block: CD16/32, CD4 BV605 RM4-5, CD8 FITC 53-6.7, CD44 APC IM7, CD69 PE-Cy7 H1.2F3, CD134 BV786 OX-86, CD137 PE 1AH2, CD45R/B220 Per-CP 5.5 RA3-6B2 (all BD Bioscience). Cells were fixed and permeabilized according to manufacturer’s protocol (BD Biosciences) and stained with aTNF*α* BV650 MP6-XT22 (BD Bioscience). Cells were washed, fixed with 2% formaldehyde and acquired on a BD LSR II and analyzed using Cytobank V7.3.0.

### T cell response assay on spleen

Mouse spleens were harvested on D105 after vaccination and single-cell suspensions were prepared by processing them through a 70-μm cell strainer (BD Biosciences). Cells were then incubated in FACS tubes at 6×10^5^ cells per tube and stimulated for 18 hours with 2 μg/ml RBD peptide mix (PepMix SARS-CoV-2 (S-RBD) Protein ID: P0DTC2 PM-WCPV-S-RBD-1, JPT) or media alone. As positive control cells were stimulated with anti-mouse CD3 [0,05 μg/ml] (Southern Biotech) and anti-mouse CD28 antibody [0,5 μg/ml] (Southern Biotech). GolgiStop (BD Biosciences) was added for the last 10 hours of the assay. Following stimulation, cells were washed, stained with Aqua Live/dead viability dye (Thermo Fisher) in PBS, washed two additional times and stained with a cocktail of monoclonal antibodies and Fc block: CD16/32, CD4 A×700 RM4-5, CD8 APC-H7 53-6.7, CD44 APC IM7 (all BD Bioscience). Cells were fixed and permeabilized according to manufacturer’s protocol (BD Biosciences) and stained for intracellular cytokines with IFN*γ* PE-Cy7 XMG1.2, TNF*α* BV650 MP6-XT22 and IL-4 BV786 11B11 (BD Bioscience). Cells were washed, fixed with 2% formaldehyde and acquired on a Cytek Aurora (Northern Lights) and analyzed using Cytobank V7.3.0.

### IFN*γ* ELISpot

The assay was performed following the manufacturer’s instructions (R&D systems, mouse IFNγ Kit Cat # EL485). In short IFNγ ELISpot analysis was performed ex vivo (without further in vitro culturing for expansion) using PBMCs depleted of CD4^+^ and enriched for CD8^+^ T cells or depleted of CD8^+^ and enriched for CD4^+^ T cells by MACS sort (Miltenyi CD4^+^ or CD8^+^ microbeads following manufacturer instructions). Tests were performed in triplicates and with a positive control (anti-CD3 monoclonal antibody (0,05 μg/ml; Southern Biotech). PVDF backed microplate plates pre-coated with IFNγ-specific antibodies (R&D systems, mouse IFNγ Kit Cat # EL485) were washed with PBS and blocked with RPMI medium (Corning) containing 5% FCS for 20min at room temperature. Per well, 5 × 10^5^ effector cells were stimulated for 16h with 2 μg/ml RBD peptide mix (PepMix SARS-CoV-2 (S-RBD) Protein ID: P0DTC2 PM-WCPV-S-RBD-1, JPT). After stimulation wells were washed and incubated with a biotinylated anti-IFNγ antibody (R&D systems, mouse IFNγ Kit Cat # EL485) overnight at 4°C. Next day wells were washed and incubated with Streptavidin-AP (R&D systems, mouse IFNγ Kit Cat # EL485) for 2h at RT. After washing wells were incubated with a 5-bromo-4-chloro-3⍰-indolyl phosphate (BCIP)/nitro blue tetrazolium (NBT) substrate (R&D systems, mouse IFNγ Kit Cat # EL485). Plates were scanned and analyzed using ImmunoSpot Microanalyzer.

### In vivo bioluminescence imaging

For bioluminescence assessment, mice were anesthetized with isoflurane gas (2% isoflurane in oxygen, 1 L/min) during injection and imaging procedures. Intraperitoneal injections of d-Luciferin (Biosynth AG) were done at a dose of 150 mg/kg, providing a saturating substrate concentration for Fluc enzyme (luciferin crosses the blood–brain barrier). Mice were imaged in a light-tight chamber using an in vivo optical imaging system (AMI HT; Spectral Instruments imaging) equipped with a cooled charge-coupled device camera. During image recording, mice inhaled isoflurane delivered via a nose cone, and their body temperature was maintained at 37°C in the dark box of the camera system. Bioluminescence images were acquired between 10 and 20 minutes after luciferin administration. Mice usually recovered from anesthesia within 2 minutes of imaging.

### White blood cell count

5 ul of blood were harvested and mixed with 45 ul 3% acetic acid with methylene blue (Stemcell) and nuclei were counted using a hematocytometer.

### Cytokine analysis

IP10, IFNa, TNFa and IL6 were measured in serum samples using the LEGENDplex™ bead-based immunoassays from Biolegend per manufacturer protocol. Assay was analyzed on a BD FACSCalibur.

### Complete blood count

CBC analysis was performed by the animal diagnostic lab at Stanford. Automated hematology was performed on an Sysmex XN-1000V analyzer system. Blood smears were made for all CBC samples and reviewed by a clinical laboratory scientist. Manual differentials were performed as indicated by species and automated analysis.

### Liver enzyme analysis

Liver enzymes were analyzed by the animal diagnostic lab at Stanford. Chemistry analysis was performed on the Siemens Dimension EXL200/LOCI analyzer. A clinical laboratory scientist performed all testing, including dilutions and repeat tests as indicated, and reviews all data.

### Safety statement

For all mouse experiments no unexpected or unusually high safety hazards were encountered

## Acknowledgments

We thank the Dynavax Corporation for the gift of the CpG formulation SD101. This work was supported by Fast Grants (to R.L), Stanford University Innovative Medicines Accelerator (IMA) (R.L., R.M.W., P.A.W.); The National Science Foundation Grant NSF CHE-2002933 (to R.M.W.); NIH R01 CA031845 (P.A.W.); The Child Health Research Institute at Stanford University and the SPARK Translational Research Program in the Stanford University School of Medicine (R.M.W., P.A.W., R.L). The support of German cancer aid (Deutsche Krebshilfe- Mildred Scheel postdoctoral fellowship to J.J.K.L) and the Stanford Cancer Translational Nanotechnology Training T32 Training Grant T32 CA196585 funded by the National Cancer Institute (T.R.B) is gratefully acknowledged. BGM is grateful for financial support from a Cancer-TNT Fellowship. Research reported in this publication was supported by the National Cancer Institute of the National Institutes of Health under Award Number F32CA254128. The content is solely the responsibility of the authors and does not necessarily represent the official views of the National Institutes of Health.

## Supporting Information

supplementary Figure S1: CART characterization, NMR spectrum, zeta potential

supplementary Figure S2: IVT mRNA quality control and immunogenicity, mRNA expression by Western Blot, upregulation of activation markers upon in vivo administration by flow cytometry

supplementary Figure S3: early RBD-specific immunoglobulin responses on D4 and 14, no induction of aHis immunoglobulins, ELISA

supplementary Figure S4: RBD mRNA + CpG-CART induces an early immunoglobulin isotype switch and generates high levels of ACE-2-RBD antibodies, ELISA and

supplementary Figure S5: Route of administration- intravenous, subcutaneous and intramuscular RBD mRNA + CpG-CART vaccine administration induces robust isotype switched anti-RBD immunoglobulin responses

supplementary Figure S6: Induction of neutralizing antibodies is independent of CpG source, RBD mRNA induced antibodies are able to bind to RBD in whole Spike protein and lead to efficient neutralization

supplementary Figure S7: aRBD antibody titers are independent of CpG source, ELISA D21 and 28

supplementary Figure S8: intracellular cytokine staining (IFN*γ*, TNF*α*, IL-4) in CD4^+^ and CD8^+^ T cells on D105

supplementary Figure S9: mouse in vivo safety studies, white blood cell count, liver enzymes, serum cytokines after prime and boost

supplementary Figure S10: mRNA-CART stability D11 at room temperature, 4°C and -20°C

## Notes

### Competing Interest Statement

The authors have declared no competing interest.

