## supporting information for "An mRNA SARS-CoV-2 vaccine employing Charge-Altering Releasable Transporters with a TLR-9 agonist induces neutralizing antibodies and T cell memory"

**SI Information**

### Supplementary Figure S1

A

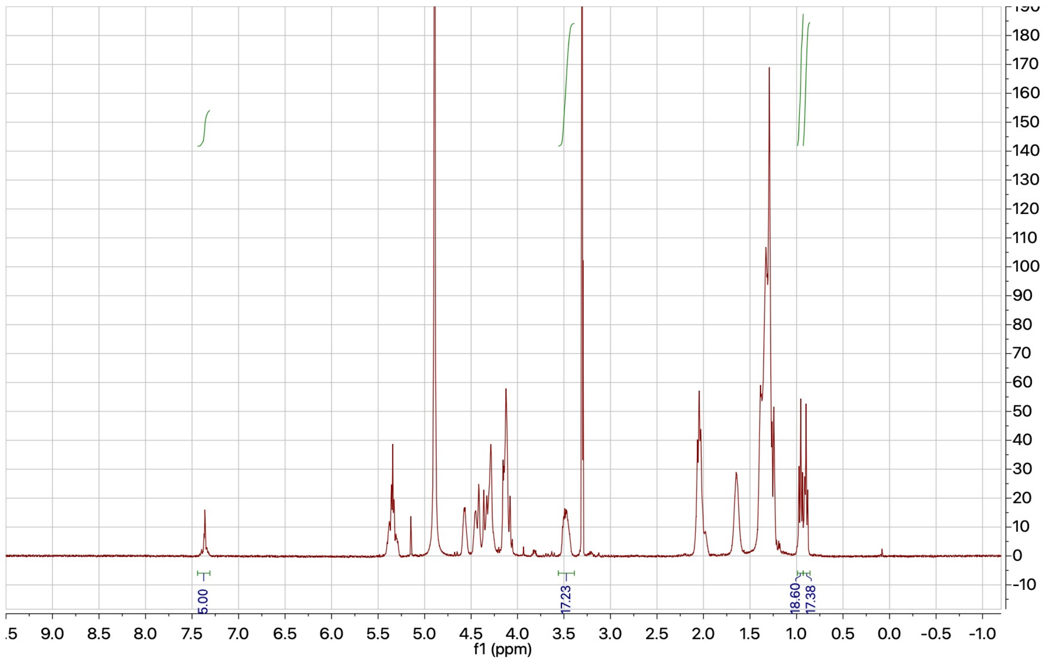

B

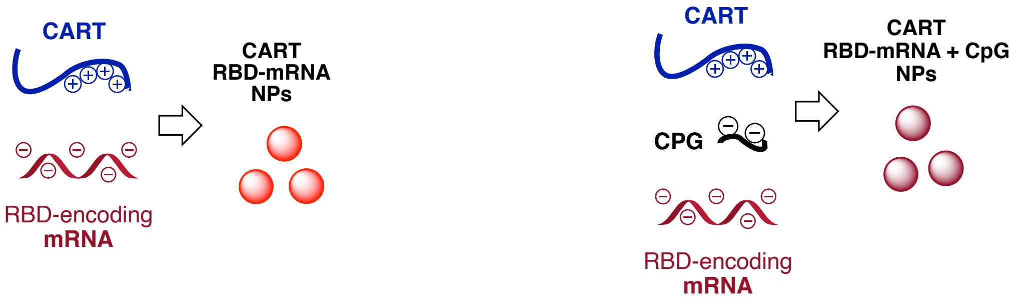


| CART/nucleotide | [a] Distribution avg size (nm) | [b] Zeta potential (mV) | [c] Encapsulation efficiency |
|---|---|---|---|
| CART RBD-mRNA | 194 +/- 41 | 41 +/- 5 | 89 % |
| CART RBD-mRNA + CpG | 327 +/- 28 | 22 +/- 9 | 91 % |

**Supplementary Figure S1) CART oligomer and CART/mRNA NP characterization A.)** $^1$H NMR spectrum ($CD_3OH$, 500 MHz) of CART $O_6$-stat-$N_6$: $A_9$. Relative ratios in $CD_3OD$: BnOH end group 7.35 ppm, 5H($C_5H_5$); nonenyl segment 0.9 ppm, 18H($CH_3$); oleyl segment 0.85 ppm, 18H($CH_3$), aminoester segment 3.5 ppm, 18H($CH_2$). Number avg. molecular weight (gel permeation chromatography vs. polystyrene standards on N-Boc protected oligomer): Mn = 4.3 kDa, D = 1.19. Comparing these ratios indicates a statistical triblock CART with DP 1:6:6:9 **B.) CART-nucleotide nanoparticle formulation and physical characterization**. [a]Nanoparticle sizes were measured using dynamic light scattering (DLS). Each value is the average from 3 trials using independently prepared formulations with the corresponding standard deviation in the same units. The plot of intensity of scattered light as a function of nanoparticle size from analysis of these samples exhibits one, monomodal distribution of particle sizes. [b]Zeta potential measurements were conducted using electrophoretic light scattering (ELS). Each value is the average of 6 measurements from 3 trials using independently prepared formulations (for each trial, 2 measurements were acquired consecutively) with the corresponding standard deviation in the same units. [c]Encapsulation efficiency was measured by the fluorescence of the RNA-specific Qubit RNA BR dye (Q10210; Invitrogen). RBD-mRNA or RBD-mRNA +CpG were complexed using a total 840 ng of nucleotide cargo and enough CART for a net 10:1 (cation:anion) ratio. This was added to 0.75 mL of RNase- free deionized water containing 5 μL of Qubit reagent. The fluorescence of the solution was immediately measuring using an excitation wavelength of 630 (4-nm slit width) and emission wavelength of 680 nm (4 nm slit width). Percent encapsulation was determined by subtracting the fluorescence of the Qubit dye alone (no nucleotide) and normalizing to the fluorescence of Qubit with uncomplexed mRNA.

### Supplementary Figure S2

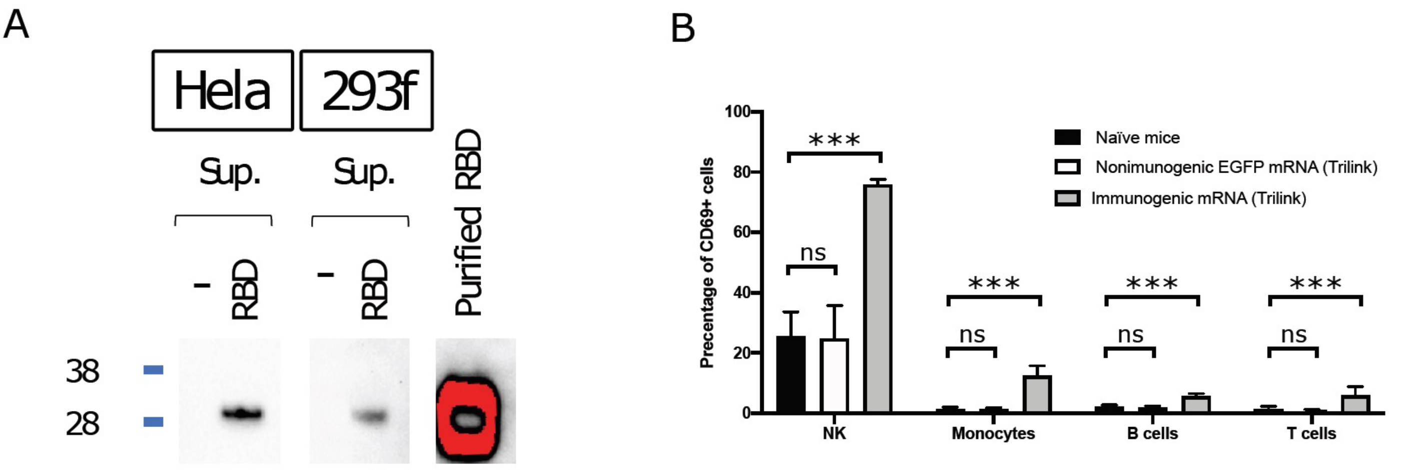


**Supplementary Figure S2) CARTs are free of nonspecific stimulation.** IVT mRNA was produced in house. Hela or 293F cells were transfected with CART-RBD mRNA. 16h later the supernatant was analyzed by western blot for RBD-His expression. Negative controls are media from non-transfected cells. 100ng of purified RBD protein was used as positive control (A). CD69 upregulation on circulating immune cell subsets 24h after IV injection of mRNA-CART complex using indicated mRNA. Non-immunostimulatory mRNA EGFP mRNA (5moU) (L-7201 Trilink), Immunostimulatory mRNA: CleanCap™ FLuc mRNA (L-7602 Trilink) (B). Puled data from multiple independent experiments. Statistical significance was assessed by One-Way ANOVA: P>0.05 (ns), P≤0.001(***).

#### Supplementary Figure S3

A

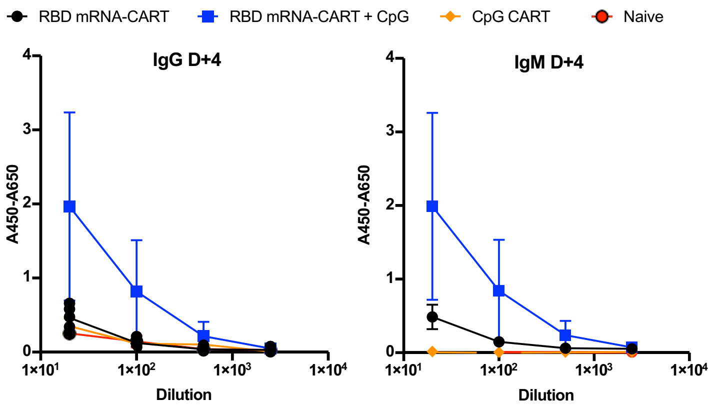


B

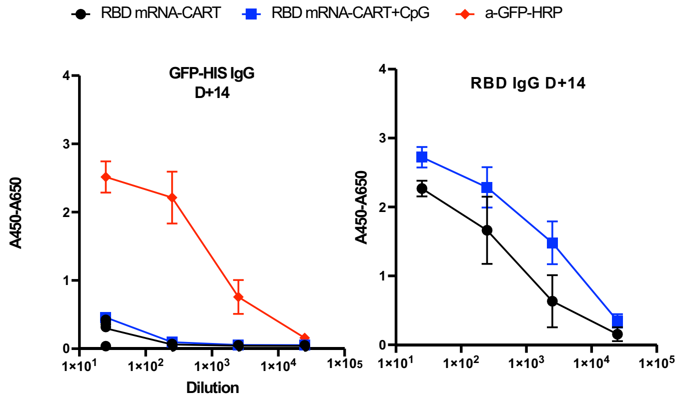


**Supplementary Figure S3) RBD mRNA + CpG-CART elicits anti-RBD immunoglobulin responses as early as 4 days post priming.** BALB/c mice were immunized intravenously with either 3ug RBD-CARTS (n=5), 3ug RBD-CART mRNA plus 3ug CpG (n=5), CpG CART (n=5), or Naïve untreated (n=5) and boosted on Day +4 and Day +8. Serum levels of RBD-specific IgGs from RBD-CART mRNA (black), RBD-CART mRNA plus CpG (blue), CpG CART (orange), and Naïve (red) mice were tested 4 days post priming by ELISA (A). The same serum samples were tested for cross reactivity with an irrelevant His-tagged GFP (GFP-His) protein using ELISA plate coated with 5ug/ml GFP-His protein and compared to an anti His antibody positive plate control (red) (B).

### Supplementary Figure S4

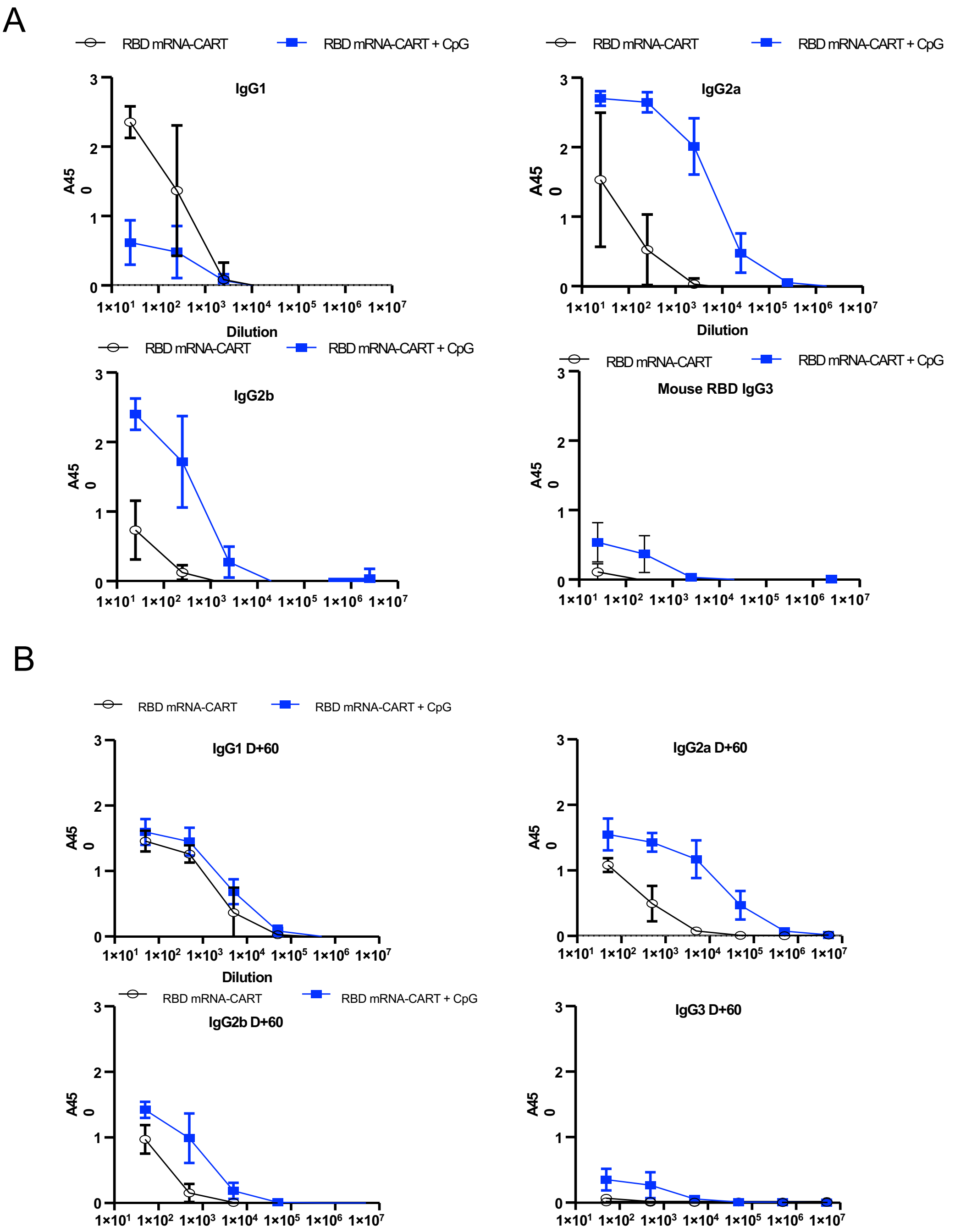


**Supplementary Figure S4) RBD mRNA + CpG-CART induces an early immunoglobulin isotype switch and generates high levels of ACE-2-RBD neutralizing antibodies.**

BALB/c mice were immunized intravenously with either 3ug RBD-CARTS (n=5), 3ug RBD-CART mRNA plus 3ug CpG (n=5), CpG CART (n=5), or Naïve untreated (n=5) and boosted on D4 and D8 after priming. Serum from RBD-CART mRNA (black), RBD-CART mRNA plus CpG (blue), immunized mice was collected on Day +14 (A) and Day +60 (B) and the distribution of IgG isotypes specific to RBD was analyzed using anti-mouse IgG1, IgG2a, IgG2b and IgG3 monoclonal antibodies in ELISA.

### Supplementary Figure S5

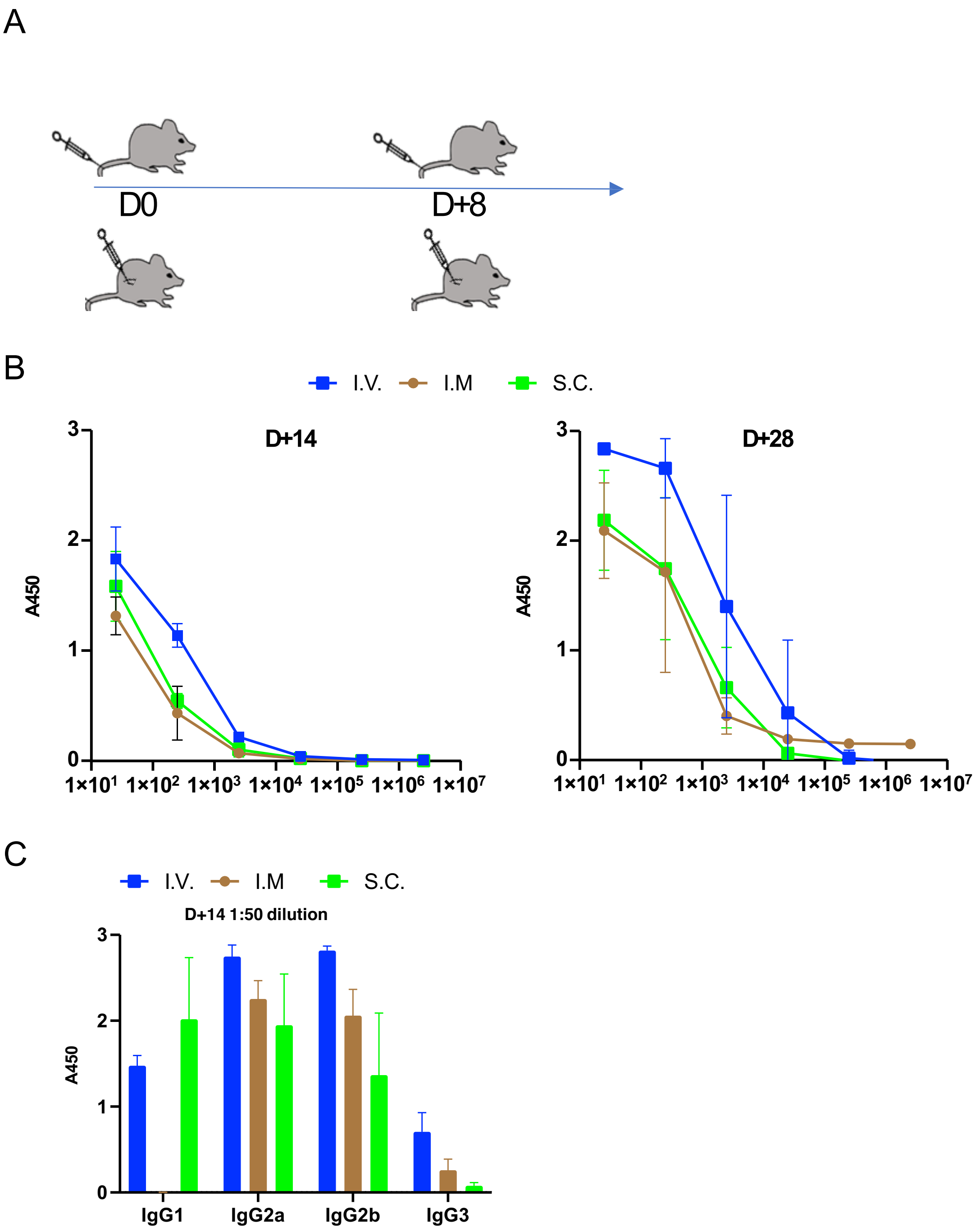


**Supplementary Figure S5) Subcutaneous and intramuscular RBD mRNA + CpG-CART vaccine administration induces robust isotype switched anti-RBD immunoglobulin responses**. 5 BALB/c mice per group were immunized intravenously (IV), intramuscular (IM), or sub cutaneous (SC) with 3ug RBD-CART mRNA plus 3ug CpG and boosted on D8 after priming (A). RBD-specific Immunoglobulin titers in serum were measured on D14 and D28 (B). RBD-specific IgG1, IgG2a, IgG2b, and IgG3 was measured on Day +14. Data representative of two individual experiments.

### Supplementary Figure S6

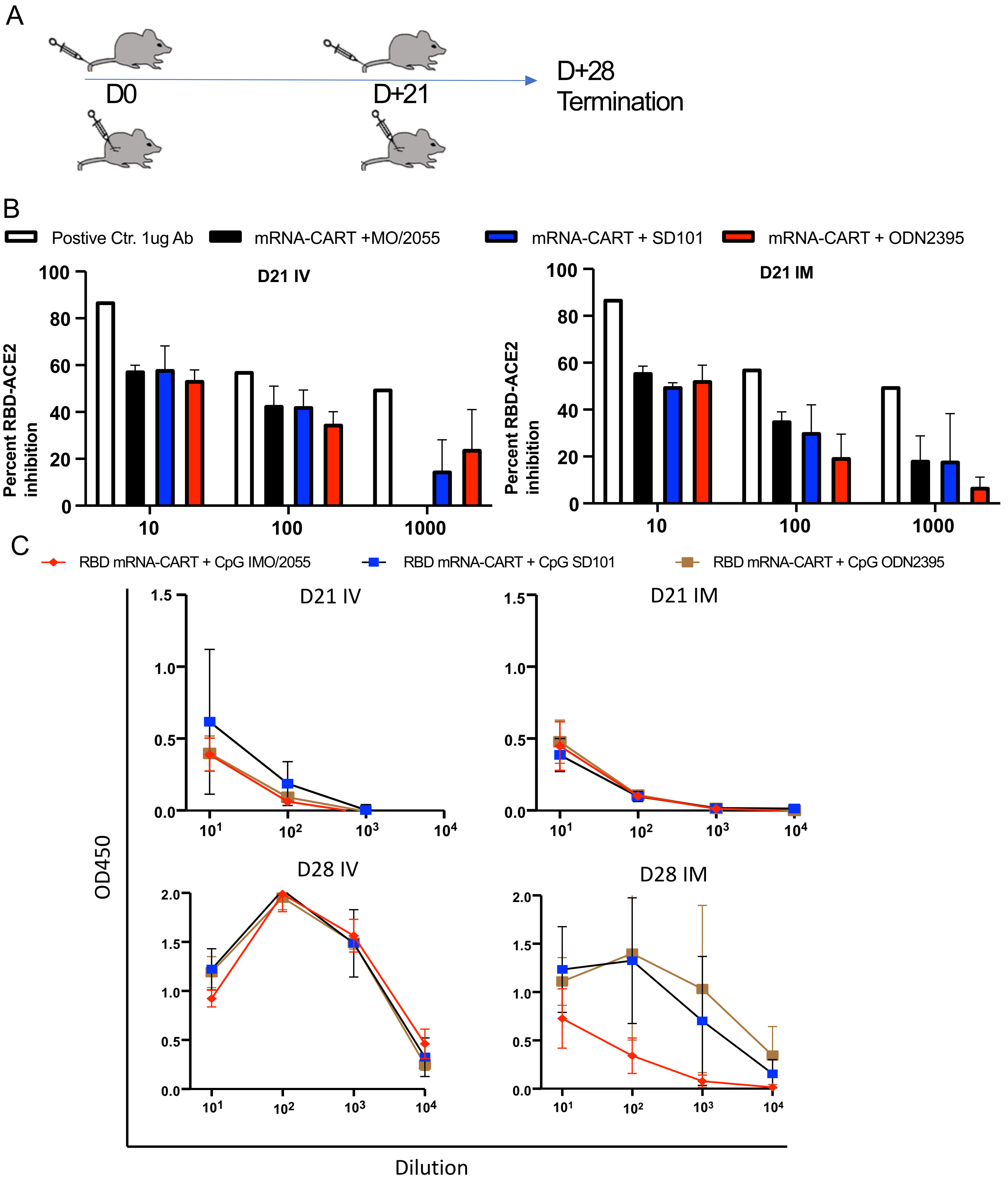


**Supplementary Figure S6) Induction of neutralizing antibodies is independent of CpG source.** 5 BALB/c mice per group were immunized intravenously (IV) or Intramuscular (IM) with 3ug RBD-CART mRNA plus 3ug CpG using 3 different sources of CpG (IMO/2055, SD101 or ODN2395) and boosted on D21 after priming (A). Serum from immunized mice harvested on D21 and D28 was tested for ability to inhibit RBD-ACE-2 binding using a commercially available RBD-ACE-2 inhibition assay (B). Serum from immunized mice harvested on D21 and D28 was tested for ability to bind to the full Spike protein by ELISA (C).

Supplementary Figure S7

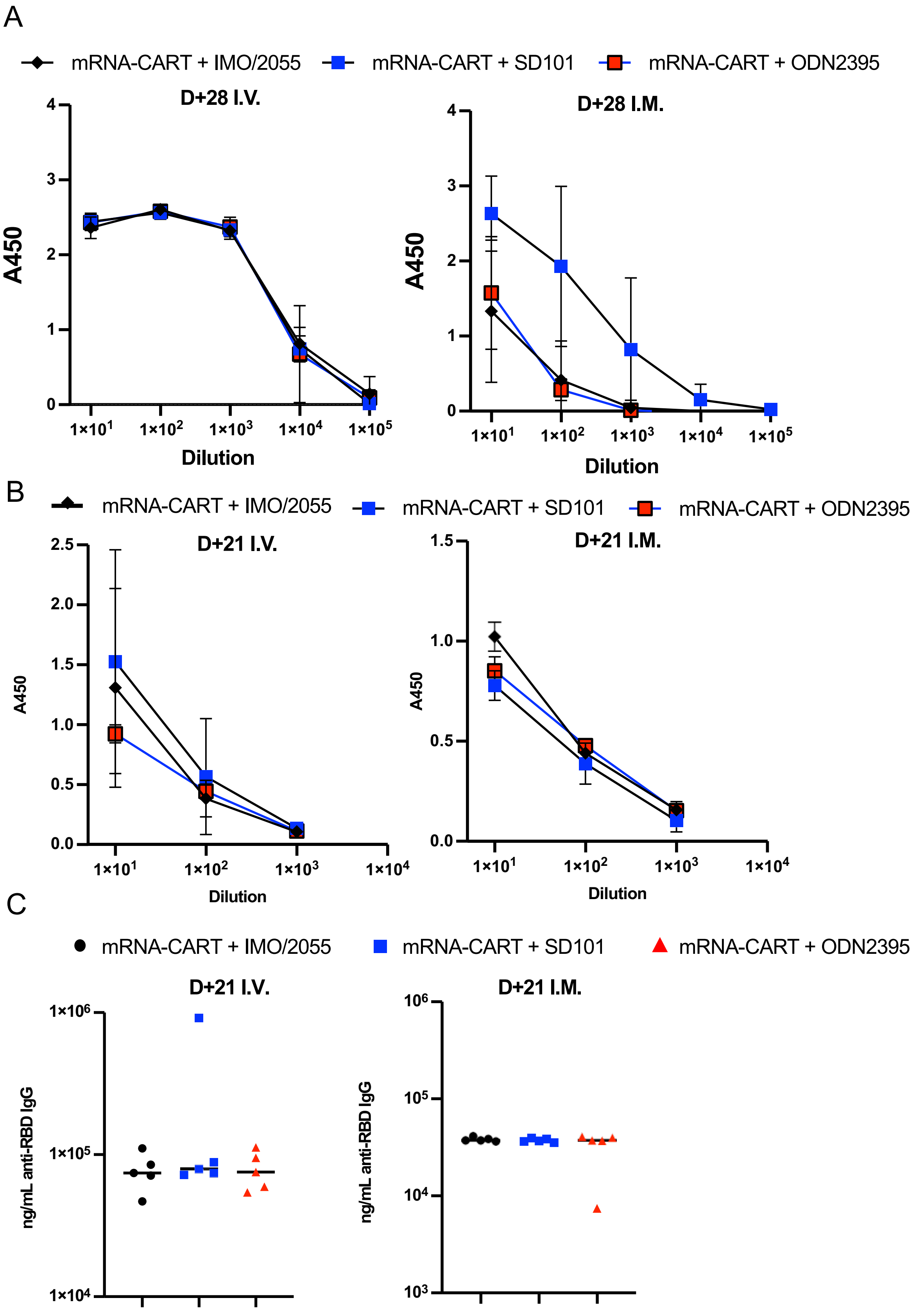


**Supplementary Figure S7) aRBD antibody titers are independent of CpG source**

Mice were immunized as described in Supplementary Fig. 6. RBD-specific Immunoglobulin titers in serum were measured on D28 (A) and D21(B). Quantification of absolute concentration of RBD-specific antibodies in serum on D21 (C).

### Supplementary Figure S8

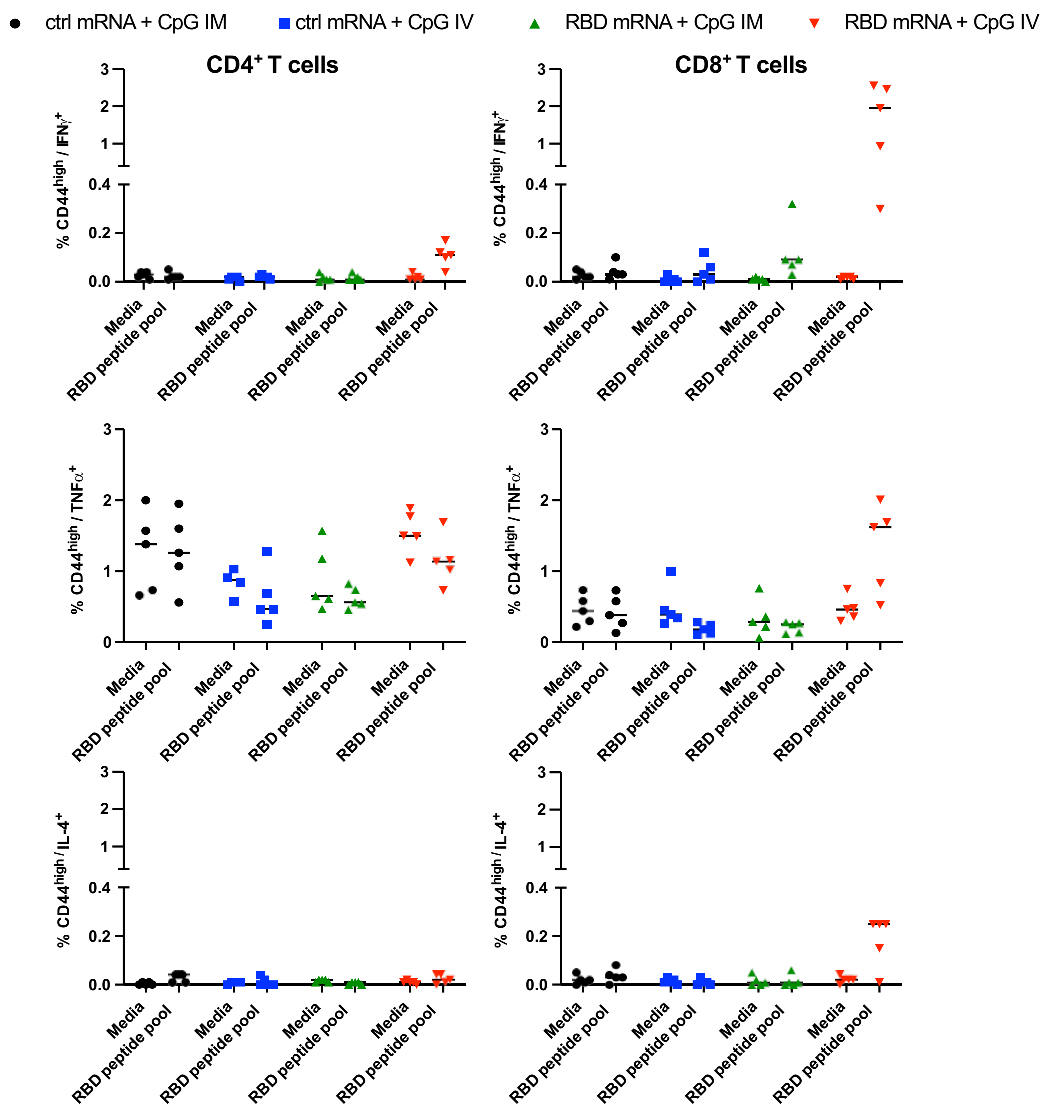


**Supplementary Figure S8)** Splenocytes from mice vaccinated with 3µg RBD-mRNA + 3µg CpG-CART or 3µg ctrl mRNA + 3µg CpG-CART either IV or IM on D1 and D21 were harvested on D105 after vaccination and incubated for 18h with RBD peptide pool or media. After stimulation cells were analyzed for IFN$\gamma$, TNF$\alpha$ or IL-4 production by intracellular cytokine staining flow cytometry.

### Supplementary Figure S9

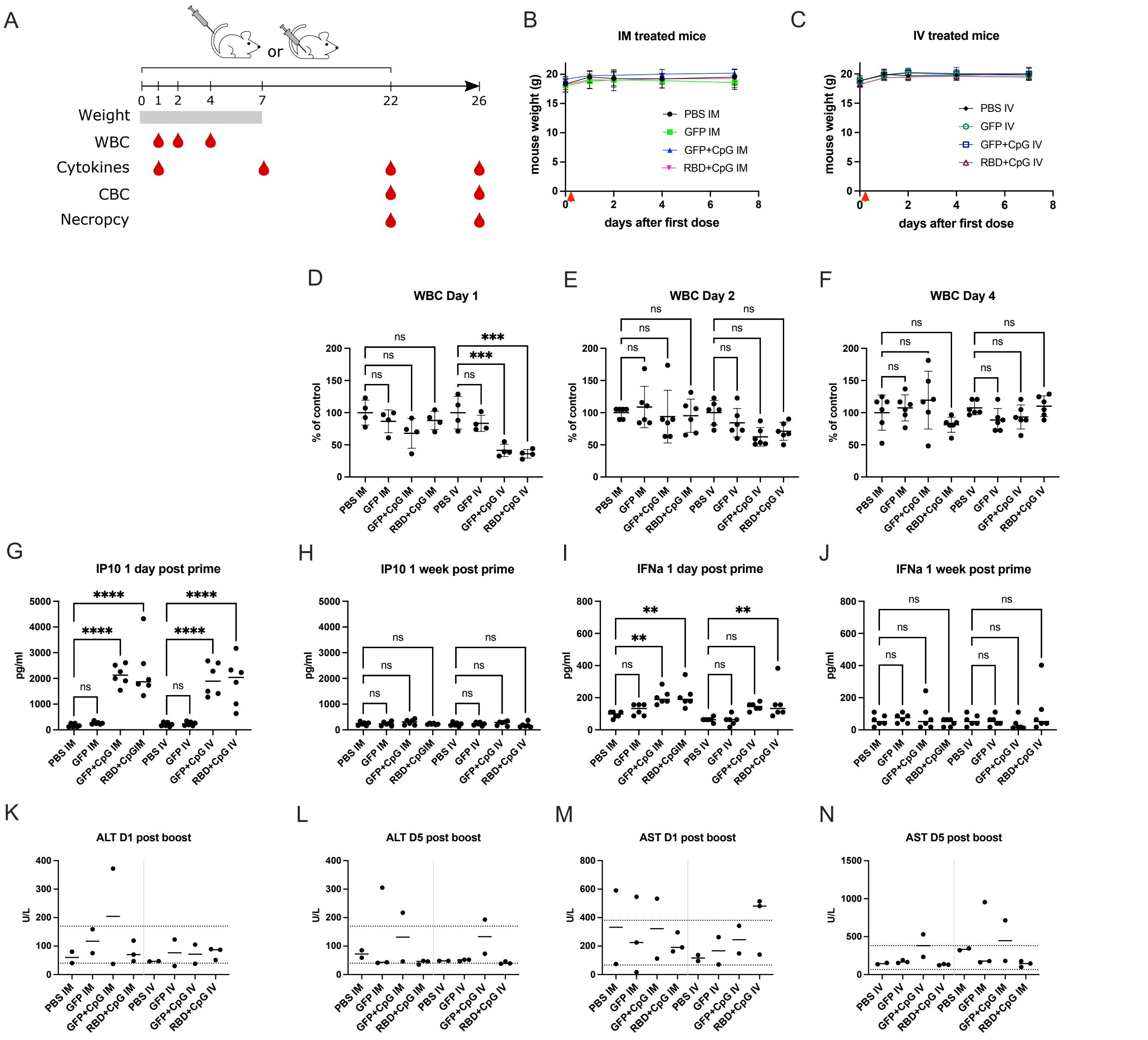


**Supplementary Figure S9: Safety evaluation of the RBD mRNA-CpG-CART Sars-Cov-2 vaccine.** Mice (6 mice per group) were injected with PBS, 3ug GFP mRNA formulated with CART or 3ug GFP mRNA + 3ug CpG formulated with CART or 3ug RBD mRNA + 3ug CpG formulated with CART. Treatments were injected in the tail vain (IV) or in the muscle (IM) on day 0 (prime) and day 21 (boost). (A) Schematic representing the schedule of injection and safety measurements. Body weight of the mice was measured on Day 0 (before first treatment), Day 1, Day 2, Day 4 and Day7 for IM (B) and IV (C) treated mice. White Blood Cell (WBC) count was assessed on Day 1 (D), Day 2 (E) and Day 4 (F). Data displayed as percentage relative to the control group. Sera were harvested on day 1 and day 7 for cytokines measurement. On Day 22 and Day 26, 3 mice per group were sacrificed and necropsy assessment and complete blood count were assessed, and serum was collected for cytokines and liver enzymes measurements. (G-J) IP10 and IFN$\alpha$ were measured in the sera harvested on Day 1 (D1 post prime) and Day 7 (1 week post prime). (K-L) Alanine transferase (ALT) and (M-N) aspartate transferase (AST) was measured in serum on Day 22 (D1 post boost) and Day 26 (D5 post boost). The experiment was performed once. Statistical significance was assessed by One-Way ANOVA: P>0.05 (ns), P≤0.05 (*), P≤0.01 (**), P≤0.001(***), P≤0.0001(****).

### Supplementary Figure S10

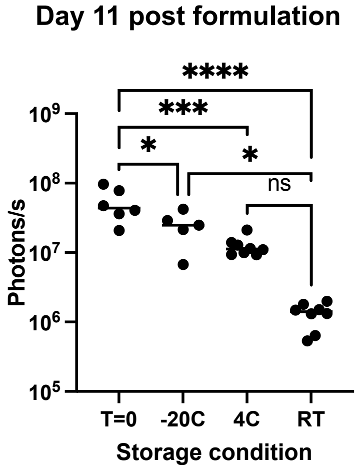


**Supplementary Figure S10: CART storage stability assay.** CARTs were formulated with Fluc mRNA at a 10:1 +/- ratio. These formulations were stored at either RT, 4C, or -20C. After storing CART formulations for 11 days the samples were thawed and injected into tail vein (50 uL total volume, 5 ug mRNA per mouse). At the same time, a fresh formulation of CART/Fluc mRNA was prepared and injected into the tail vein. The levels of luciferase expression were determined by BLI (units: p/s), comparing efficacy to the freshly formulated CART/mRNA. The experiment was performed once. Statistical significance was assessed by One Way Anova: $P>0.05$ (ns), $P\leq0.05$ (*), $P\leq0.01$ (**), $P\leq0.001$(***), $P\leq0.0001$(****).
